# VTA μ–opioidergic neurons facilitate low sociability in protracted opioid withdrawal

**DOI:** 10.1101/2024.07.08.602522

**Authors:** Adrienne Y. Jo, Yihan Xie, Amrith Rodrigues, Raquel Adaia Sandoval Ortega, Kate Townsend Creasy, Kevin T. Beier, Julie A. Blendy, Gregory Corder

## Abstract

Opioids initiate dynamic maladaptation in brain reward and affect circuits that occur throughout chronic exposure and withdrawal that persist beyond cessation. Protracted withdrawal is characterized by negative affective behaviors such as heightened anxiety, irritability, dysphoria, and anhedonia, which pose a significant risk factor for relapse. While the ventral tegmental area (VTA) and mu-opioid receptors (MORs) are critical for opioid reinforcement, the specific contributions of VTA^MOR^ neurons in mediating protracted withdrawal-induced negative affect is not fully understood. In our study, we elucidate the role of VTA^MOR^ neurons in mediating negative affect and altered brain-wide neuronal activities following opioid exposure and withdrawal in male and female mice. Utilizing a chronic oral morphine administration model, we observe increased social deficit, anxiety-related, and despair-like behaviors during protracted withdrawal. VTA^MOR^ neurons show heightened neuronal FOS activation at the onset of withdrawal and connect to an array of brain regions that mediate reward and affective processes. Viral re-expression of MORs selectively within the VTA of MOR knockout mice demonstrates that the disrupted social interaction observed during protracted withdrawal is facilitated by this neural population, without affecting other protracted withdrawal behaviors. Lastly, VTA^MORs^ contribute to heightened neuronal FOS activation in the anterior cingulate cortex (ACC) in response to an acute morphine challenge, suggesting their unique role in modulating ACC-specific neuronal activity. These findings identify VTA^MOR^ neurons as critical modulators of low sociability during protracted withdrawal and highlight their potential as a mechanistic target to alleviate negative affective behaviors associated with opioid withdrawal.

**SIGNFICANCE:** The compelling urge for relief from negative affective states during long-term opioid withdrawal presents a crucial challenge for maintaining abstinence. The ventral tegmental area (VTA) and its mu-opioid receptor-expressing (VTA^MOR^) neurons represent a critical target of opioidergic action that underlie dependence and withdrawal. Chronic activation of VTA^MOR^ neurons during opioid exposure induces maladaptations within these neurons and their structurally connected circuitries, which alter reward processing and contribute to negative affect. Using an oral morphine drinking paradigm to induce dependence, we demonstrate that withdrawal engages VTA^MOR^ neurons and identify this neuronal population as key mediators of opioid withdrawal-induced social deficits. These findings hold promise to inform development of targeted therapies aimed at alleviating negative affective states associated with protracted opioid withdrawal.

## INTRODUCTION

Protracted opioid withdrawal—the weeks-to-months long period after drug cessation—is marked by negative affective behaviors such as heightened anxiety, irritability, dysphoria, and anhedonia, which continue long after acute physical withdrawal symptoms have subsided. The compelling urge to relieve this negative state poses a significant challenge for sustained abstinence, with relapse rates among individuals with opioid use disorder (OUD) exceeding 90% (Weiss, 2011; Nwaefuna, 2023). Pre-clinical models of protracted withdrawal using primarily investigator-administered opioids have extensively assessed the enduring effects of negative affect during spontaneous withdrawal. These effects, including increased anxiety- and despair-related behaviors, anhedonia, and social deficits, persist long after the last opioid administration (Goeldner et al., 2011; Bravo et al., 2020; Welsch et al., 2020; Becker et al., 2021; Pomrenze et al., 2022; Ozdemir et al., 2023). These observations underscore the critical need to investigate the underlying neuroadaptations resulting from repeated opioid exposure and withdrawal that facilitate the development of these maladaptive behaviors (Koob and Volkow, 2010, 2016).

Central to the brain’s reward circuitry, the ventral tegmental area (VTA) is a critical site for opioid action and reinforcement, and is enriched with mu-opioid receptor (MOR) – containing neurons. MORs are essential for opioid-induced analgesia, nociception, and dependence (Matthes et al., 1996), and their activation, specifically in the VTA, is necessary for opioid-induced reward (Phillips and LePiane, 1980; Britt and Wise, 1983; Olmstead and Franklin, 1997; Fields and Margolis, 2015) and the reinstatement of drug-seeking behavior (Stewart, 1984). Beyond their well-established role in opioidergic activity and reward processing, MORs have been proposed to play a fundamental role in mediating social behaviors, with previous studies demonstrating MORs are critical for social interaction (Becker et al., 2014; Pellissier et al., 2018; Toddes et al., 2021).

Although various regions such as the nucleus accumbens (Pomrenze et al., 2022; Fox et al., 2023), central amygdala (Jiang et al., 2021), bed nucleus of the stria terminalis (Delfs et al., 2000), habenula (Valentinova et al., 2019; Bailly et al., 2023), and dorsal raphe nucleus (Lutz et al., 2014; Pomrenze et al., 2022; Welsch et al., 2023) have been studied for their role in negative affect circuitry, the specific contributions of VTA^MOR^ neurons to behavioral effects during protracted withdrawal remain elusive. Characterized as a central hub within the mesocorticolimbic circuitry, the VTA’s extensive afferent and efferent projection targets (Bouarab et al., 2019) position the region to influence a wide range of behaviors related not only to reward, but also to stress and aversion. These unique structural connections, alongside the interplay of opioid action on VTA^MOR^ neurons during morphine administration, suggest a fundamental role for the VTA in the emergence of maladaptive behaviors associated with repeated opioid use and withdrawal.

Here, we aimed to elucidate the contribution of VTA^MOR^ neurons in the development of morphine dependence and negative affect during protracted withdrawal. We utilized a chronic oral morphine drinking paradigm followed by protracted withdrawal and conducted a targeted examination of VTA^MOR^ neurons using MOR-specific mouse lines and Cre-dependent viral tools. Leveraging the genetic neural activity marker, c-FOS, we show that VTA^MOR^ neurons across the anterior-posterior axis are activated during dependence and naloxone-precipitated withdrawal, indicating a neural adaptation within this cell type that may be in-volved in the long-lasting behavioral effects observed during protracted withdrawal. Additionally, using viral antero- and retrograde tracing to map the input-output connectivity architecture of VTA^MOR^ neurons, we discovered a diverse array of brain-wide connections with regions implicated in stress and negative affect, including the prefrontal cortex and locus coeruleus. Notably, we demonstrated that VTA^MOR^ neurons mediate reduced sociability during protracted withdrawal through viral MOR re-expression within the VTA of MOR KO mice. Furthermore, these neurons contribute to increased FOS expression in the anterior cingulate cortex (ACC) following an acute morphine challenge injection, an effect that is diminished in mice with a prior history of morphine exposure. In total, our results highlight the critical role of VTA^MOR^ neurons in facilitating low sociability during protracted opioid withdrawal and propose this neuronal population as a potential target for interventions aimed at alleviating the distressing symptoms associated with protracted withdrawal.

## RESULTS

### Protracted morphine withdrawal leads to the emergence of negative affective behaviors

To establish a model of morphine dependence and negative affective behaviors during protracted withdrawal, we employed a chronic, forced oral morphine drinking paradigm with escalating concentrations over 13 days, followed by 4 weeks of protracted withdrawal, and a behavioral battery [social interaction test, elevated zero maze, tail suspension test, and hot plate test] **(Fig. 1a)**. Morphine-treated mice show an increased volume consumption of morphine relative to day 1 throughout the paradigm **(Fig. 1b)**, and exhibit escalated morphine dose intake throughout the 13-day exposure period **(Fig. 1c)**. 24 hours following the cessation of morphine administration, mice undergoing spontaneous withdrawal demonstrate physical dependence, as shown through an increased global withdrawal score relative to opioid-naïve controls **(Fig. 1d;** unpaired t test, t=8.014, df=27, p<0.0001). This result is driven by increased frequencies of paw tremors, head shakes, body shakes, and genital licking and grooming bouts **(Extended Data Fig. 1-1b-e)**.

**Figure 1:**
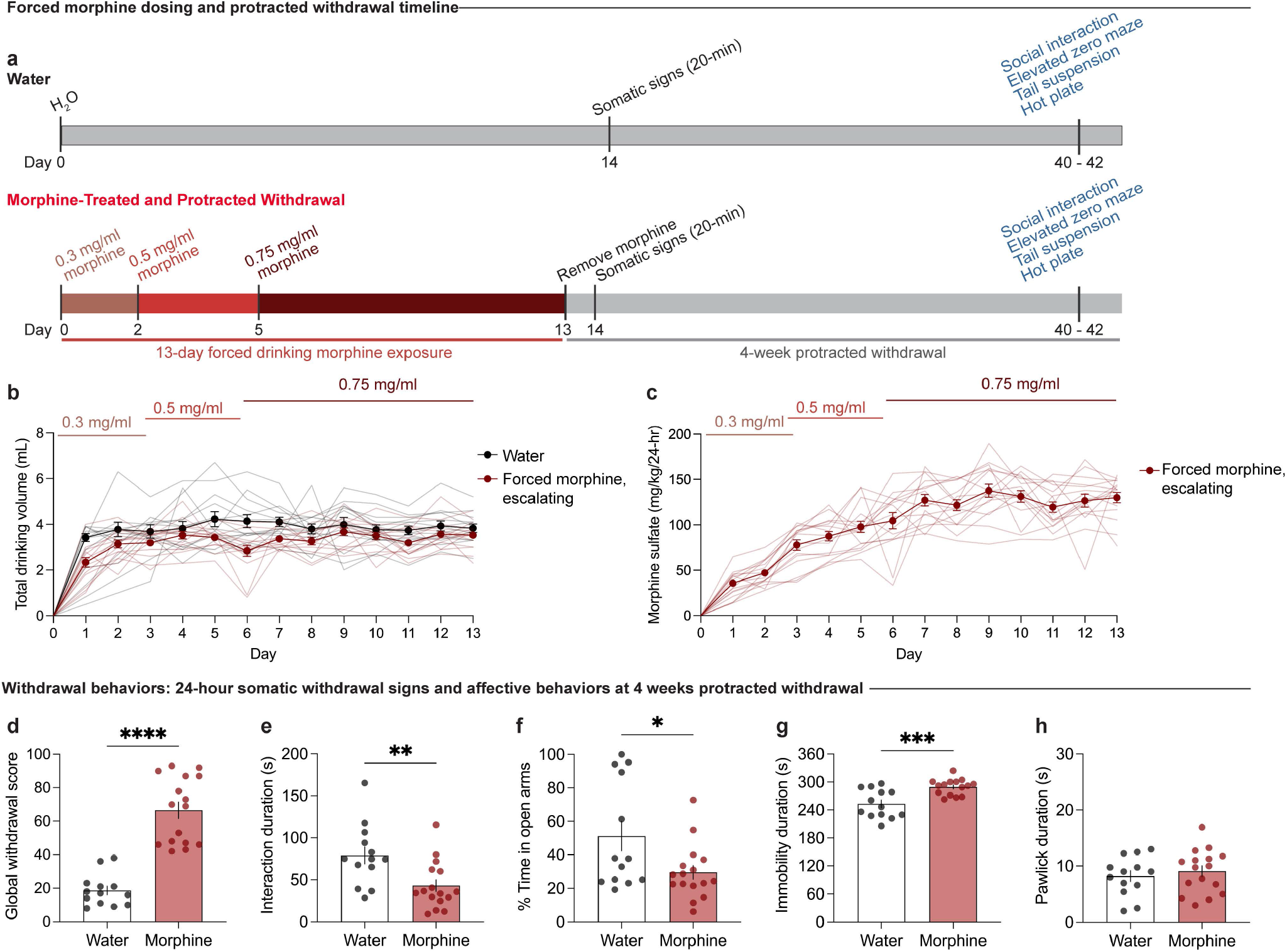
Protracted morphine withdrawal leads to the emergence of negative affective behaviors. **(a)** Experimental schematic of the 13-day chronic forced morphine drinking exposure with escalating concentrations, assessment of somatic withdrawal signs 24-hours following morphine cessation, a 4-week protracted withdrawal period, and behavioral battery. **(b)** Daily consumed volumes in the opioid naïve and morphine-treated groups across the 13-day drinking paradigm. **(c)** Daily morphine sulfate dose administration across the 13-day drinking exposure period. **(d)** At 24 hours following cessation, morphine-treated mice undergoing spontaneous withdrawal exhibit an increased global withdrawal score (n=13 opioid naïve/16 morphine-treated, unpaired t test, t=8.014, df=27, p<0.0001). At 4-weeks of protracted morphine withdrawal, mice display **(e)** low sociability as assessed through the social interaction test (unpaired t test, t=2.974, df=27, p=0.0061), **(f)** anxiety-like behavior as assessed through the elevated zero maze assay (unpaired t test, t=2.357, df=27, p=0.0259), **(g)** increased immobility as evaluated through the tail suspension test (unpaired t test, t=3.937, df=26, p=0.0006), and **(h)** unaltered nociceptive hypersensitivity as assessed through the hot plate test (unpaired t test, t=0.6185, df=27, p=0.5414). Data = mean ± SEM, *p<0.05, **p<0.01, ***p<0.001, ****p<0.0001.

Following 4 weeks of protracted morphine withdrawal, morphine-treated mice demonstrate social deficit-like behavior in the social interaction test, as demonstrated through a decreased social interaction duration **(Fig. 1e;** unpaired t test, t=2.974, df=27, p=0.0061). Morphine-treated mice also exhibited increased anxiety-like behavior in the elevated zero maze, as shown through decreased time spent in the open arms **(Fig. 1f;** unpaired t test, t=2.357, df=27, p=0.0259), as well as increased despair-related behavior through an increased duration of immobility time in the tail suspension test **(Fig. 1g;** unpaired t test, t=3.937, df=26, p=0.0006). Opioid naïve and morphine-treated mice did not show differences in the total paw lick duration in the hot plate test **(Fig. 1h;** unpaired t test, t=0.6185, df=27, p=0.5414), indicating a lack of persistent alterations in nociceptive hypersensitivity during protracted morphine withdrawal.

### VTA^MOR^ neurons show increased neural activation during withdrawal in morphine-dependent mice

Having observed the emergence of negative affective behaviors, including reduced social interactions, during protracted morphine withdrawal, we next sought to determine if VTA^MOR^ neurons are engaged during dependence. Given the prominent role of the VTA in opioid withdrawal and social reward and dysfunction, (Kaufling and Aston-Jones, 2015; Bariselli et al., 2016; Hung et al., 2017; Langlois and Nugent, 2017; Porcelli et al., 2019; Solié et al., 2022), we aimed to assess the withdrawal-related activation of VTA^MOR^ neurons, as these opioid receptor-expressing cell-types would be directly engaged throughout morphine exposure and thus subject to intracellular adaptations leading to dependence. Identifying the anatomical location of precipitated withdrawal-active VTA^MOR^ neurons is key to understanding their role in the early stages of dependence, which may inform our understanding of the neural mechanisms contributing to the enduring effects observed during protracted withdrawal. Given the low reported expression of c-FOS during spontaneous withdrawal (removal of morphine drinking access), we used the MOR antagonist naloxone to precipitate withdrawal and provide a salient stimulus for induction of the immediate early gene, c-FOS. This precipitated approach allowed us to localize the activation of specific neural populations in the VTA during the critical window during the transition from acute to protracted morphine withdrawal.

To determine if MOR-expressing neurons exhibit differential neuronal activation during morphine withdrawal, we employed a targeted adeno-associated viral (AAV) approach to label MOR-containing neurons, which allowed us to pinpoint the activation of c-FOS in these neurons. A retro-orbital injection of AAV.PHP.eB*mMORp*-eYFP (a synthetic MOR promoter driving enhanced yellow fluorescent protein (eYFP) (Salimando et al., 2023) was administered prior to initiating the 13-day chronic morphine exposure paradigm that culminated in naloxone-precipitated withdrawal and immediate assessment of somatic withdrawal signs **(Fig. 2a, c)**. Mice exposed to the chronic morphine drinking paradigm with escalating concentrations show a heightened global withdrawal score following naloxone-precipitated (1 mg/kg, s.c.) withdrawal, compared to opioid naïve control mice **(Fig. 2b;** unpaired t test, t=14.14, df=6, p<0.0001), which confirmed the presence of physical dependence to morphine. The increased global withdrawal score was primarily driven increased frequencies of diarrhea and jumps **(Extended Data Fig. 2-1h-i)**.

**Figure 2:**
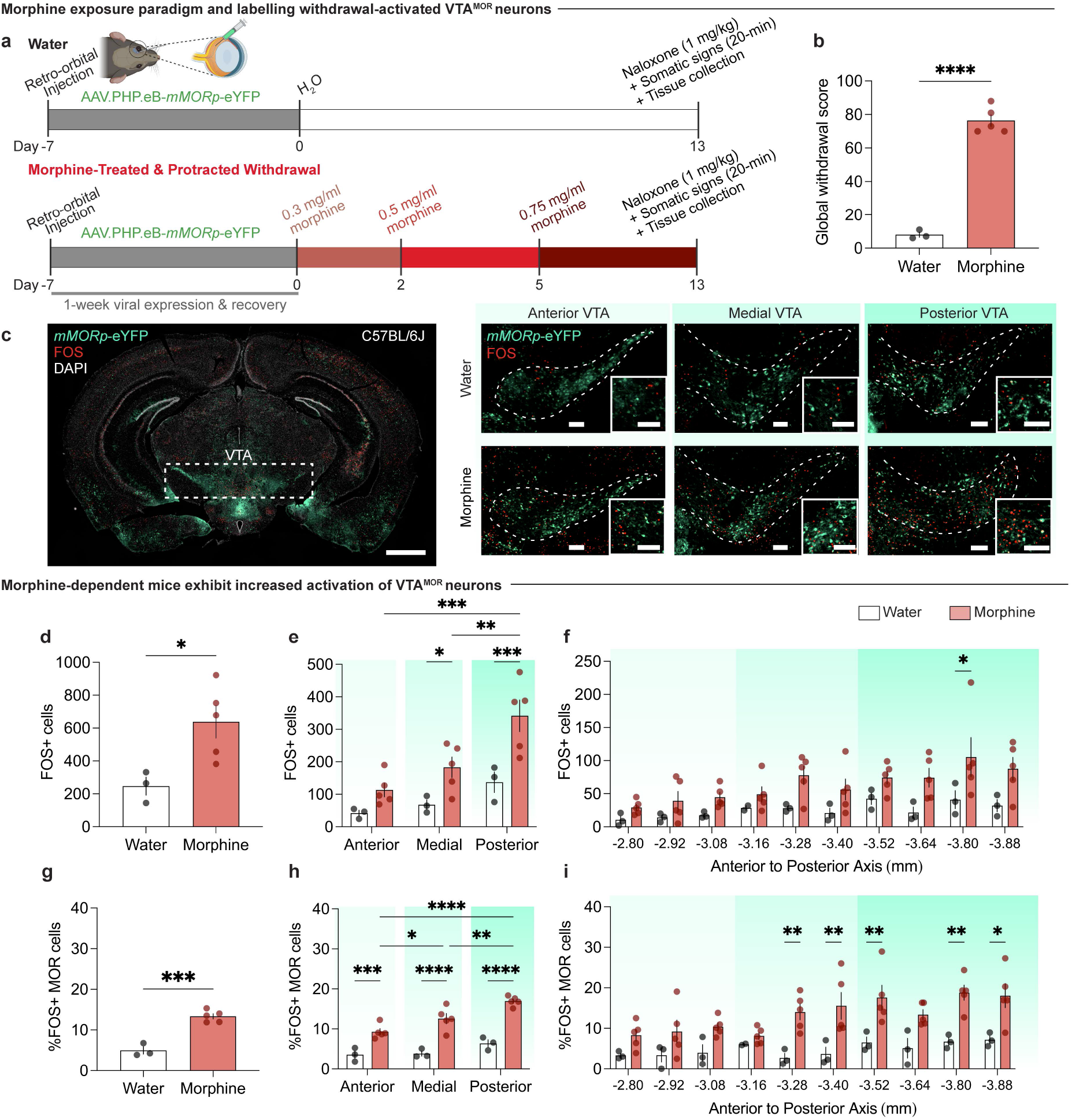
VTA^MOR^ neurons show increased neural activation during withdrawal in morphine-dependent mice. **(a)** Experimental schematic of a retro-orbital injection of AAV.PHP.eB-*mMORp*-eYFP, followed by 13-day chronic morphine drinking with escalating concentrations, naloxone-precipitated withdrawal, assessment of somatic withdrawal signs, and tissue collection 90 minutes post-naloxone injection. **(b)** Morphine-treated mice undergoing withdrawal demonstrate an increased global withdrawal score relative to opioid naïve mice (n=3 opioid-naïve/5 morphine-treated, unpaired t test, t=14.14, df=6, p<0.0001). **(c)** Coronal sections showing *mMORp*-eYFP and FOS staining (left, VTA (dashed white box highlights VTA), scale, 1 mm) in the anterior, medial, and posterior VTA in opioid naïve and morphine-treated conditions (right, scale, 200 μm). **(d)** Precipitated withdrawal-induced FOS in the VTA shows increased FOS expression in morphine-dependent mice (unpaired t test, t=2.849, df=6, p=0.0292), an effect observed in the **(e)** medial and posterior VTA (two-way ANOVA + Bonferroni, subregion F_2,18_=10.96, p=0.0008; treatment F_1,18_=20.27, p=0.0003; interaction (subregion x treatment) F_2,18_=1.834, p=0.1884). **(f)** FOS quantifications across varying bregma levels of the VTA show increased FOS expression at −3.80 mm from bregma in morphine-treated mice relative to opioid-naïve counterparts (two-way ANOVA + Bonferroni, bregma F_9,59_=2.476, p=0.0180; treatment F_1,59_=31.47, p<0.0001; interaction (bregma x treatment) F_9,59_=0.5615, p=0.8228). **(g)** Morphine-dependent mice exhibit an increased percentage of FOS+ VTA^MOR^ neurons (unpaired t test, t=7.661, df=6, p=0.0003), an effect observed in the **(h)** anterior, medial, and posterior VTA (two-way ANOVA + Bonferroni, subregion F_2,18_=14.71, p=0.0002; treatment F_1,18_=108.6, p<0.0001; interaction (subregion x treatment) F_2,18_=3.100, p=0.0697). **(i)** The percentage FOS+ VTA^MOR^ neurons across varying bregma levels shows an increased percentage at −3.28 mm, −3.40 mm, −3.52 mm, −3.80 mm, and −3.88 mm from bregma in morphine-dependent mice (two-way ANOVA + Bonferroni, bregma F_9,59_=2.654, p=0.0118; treatment F_1,59_=68.84, p<0.0001; interaction (region x treatment) F_9,59_=1.092, p=0.3828). Data = mean ± SEM, *p<0.05, **p<0.01, ***p<0.001, ****p<0.0001.

We first evaluated whether the VTA shows differential withdrawal-induced neuronal activation **(Fig. 2d-f)**. We observed an increase in naloxone-precipitated FOS expression across the anterior-posterior axis of the VTA in morphine-treated mice relative to opioid naïve mice **(Fig. 2d;** unpaired t test, t=2.849, df=6, p=0.0292). Interestingly, a significant elevation in withdrawal-induced FOS was observed in the medial and posterior VTA relative to the anterior subregion **(Fig. 2e;** two-way ANOVA + Bonferroni’s multiple comparison, subregion F_2,18_=10.96, p=0.0008; treatment F_1,18_=20.27, p=0.0003; interaction (subregion x treatment) F_2,18_=1.834, p=0.1884). Indeed, a higher percentage of withdrawal FOS+ MOR+ neurons was observed throughout the VTA **(Fig. 2g;** unpaired t test, t=7.661, df=6, p=0.0003) across the anterior, medial, and posterior VTA in morphine-treated mice **(Fig. 2h;** two-way ANOVA + Bonferroni’s multiple comparison, subregion F_2,18_=14.71, p=0.0002; treatment F_1,18_=108.6, p<0.0001; interaction (subregion x treatment) F_2,18_=3.100, p=0.0697). This finding highlights the altered activation of VTA^MOR^ neurons following the development of dependence and at the initiation of withdrawal, suggesting that this neural population undergoes adaptive processes implicated in withdrawal, which may contribute to the negative affective behavioral effects observed during protracted withdrawal.

### Structural input-output mapping of VTA^MOR^ neurons

To further investigate the circuitry associated with the activation of VTA^MOR^ neurons during withdrawal, we mapped the structural connectivity of this VTA cell-type. Identifying the direct input and output structural connections to and from VTA^MOR^ neurons can establish the distribution architecture of withdrawal-related neurotransmissions during early and protracted morphine withdrawal. We first used *Oprm1*^MOR-T2A-Cre^ mice and Cre-dependent AAV vectors for retrograde and anterograde brain-wide tracing. While the structural circuit architecture of dopaminergic and GA-BAergic neurons in the VTA is known (Sesack and Grace, 2010; Russo and Nestler, 2013; Beier et al., 2015), the specific afferent and efferent projections of VTA^MOR^ neurons have not been directly explored. Using a modified rabies virus for monosynaptic input labelling in *Oprm1*^MOR-T2A-Cre^ mice **(Fig. 3a)**, we injected Cre-dependent helper viruses and the rabies virus in the VTA **(Fig. 3b)**. We found that VTA^MOR^ neurons receive a diverse range of monosynaptic inputs **(Fig. 3c-d, Extended Data Fig. 3-1b, and Table 3)** from regions critical to reward and aversion processing, such as the striatum (NAcC, NAcSh, DLS, DMS, VLS, VMS), pallidum (VP, GP, BNST), amygdala (CeA), thalamic nuclei (MD, LHb, MHb), hypothalamus (LH, ZI, POA), midbrain structures (PAG, DRN), and the pons (PBN).

**Table 3:**
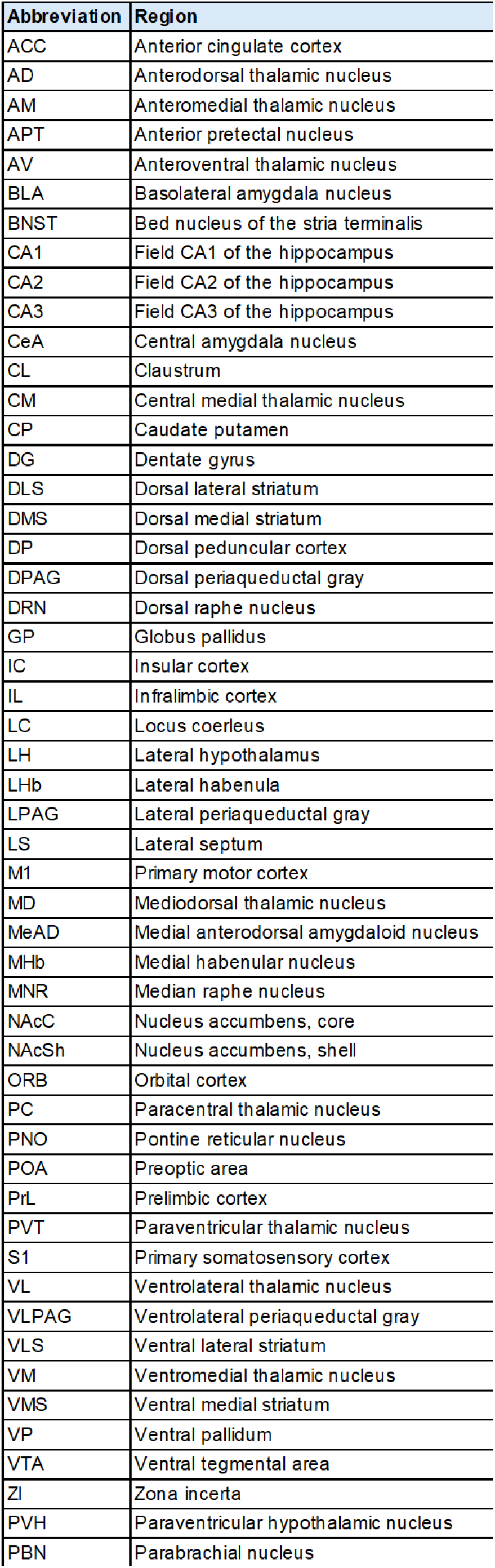
Abbreviations of brain regions.

**Figure 3:**
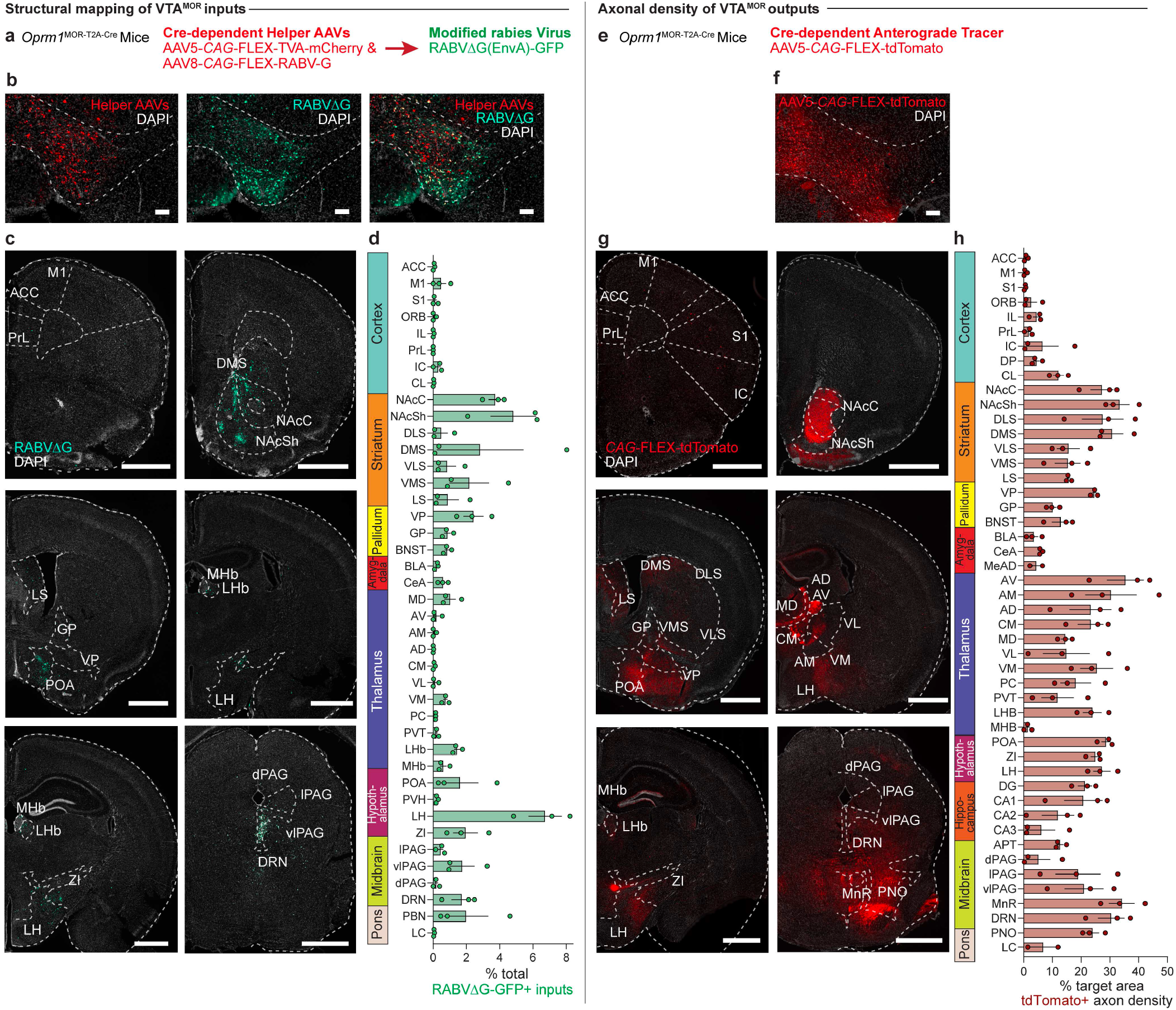
Structural input-output mapping of VTA^MOR^ neurons. **(a)** Experimental schematic for rabies-mediated monosynaptic labelling of VTA^MOR^ inputs in *Oprm1*^MOR-T2A-Cre^ mice through a unilateral injection of AAVs expressing Cre-dependent TVA-mCherry (AAV5-*CAG*-FLEX-TVA-mCherry) and rabies glycoprotein (AAV8-*CAG*-FLEX-RABV-G) helper viruses in the VTA, followed by a subsequent injection of the EnvA-pseudotyped, G-deleted, GFP-expressing rabies virus (RABVΔG-GFP) into the same VTA injection site. **(b)** Unilateral VTA injection site in *Oprm1*^MOR-T2A-Cre^ mice (scale, 200 μm). **(c)** Rabies-infected GFP-positive VTA^MOR^ inputs (scale, 1000 μm). **(d)** Quantification of VTA^MOR^ inputs presented as the fraction of GFP-positive counts per region by the total sum of all rabies-labelled inputs (n=3 *Oprm1*^MOR-T2A-Cre^ mice). **(e)** Anterograde axon tracing through a unilateral viral injection of AAV5-FLEX-tdTomato in the VTA of *Oprm1*^MOR-T2A-Cre^ mice. **(f)** Representative image of the unilateral VTA injection site of the Cre-dependent anterograde tracer (scale, 200 μm). **(g)** Coronal images of mCherry-positive VTA^MOR^ axon density projections (scale, 1000 μm). **(h)** Quantification of VTA^MOR^ axon density outputs (n=3 *Oprm1*^MOR-T2A-Cre^ mice). Data = mean ± SEM.

We also mapped the axonal projections of VTA^MOR^ outputs through anterograde tracing also in *Oprm1*^MOR-T2A-Cre^ mice **(Fig. 3e)** through an injection of a Cre-dependent anterograde tracer into the VTA **(Fig. 3f)**. Axonal density analyses show VTA^MOR^ neurons project densely to the striatum (NAcC, NAcSh, DLS, DMS, VLS, VMS), pallidum (VP, GP, BNST), the amygdala (CeA), thalamic nuclei (AV, AM, AD, CM, MD, VL, VM, LHb), hypothalamus (ZI, LH, POA), hippocampus (DG, CA1-3), midbrain (PAG, DRN), and pons (LC) (**Fig. 3g-h, Extended Data Fig. 3-1d, and Table 3)**. Together, these extensive and diverse structural connections of VTA^MOR^ neurons with key regions involved in processing reward and aversion (Russo and Nestler, 2013) suggest how these neurons may facilitate the development of negative affect throughout dependence and withdrawal, a process likely involving disrupted reward processing (Koob and Volkow, 2016).

### Morphine withdrawal leads to decreased neuronal activation following an acute morphine challenge in protracted withdrawal in the ACC and LC

From the comprehensive, brain-wide structural connectivity map of VTA^MOR^ neurons, we next aimed to understand how these anatomically connected regions, as well as additional structures implicated in withdrawal, respond to an acute morphine challenge injection in mice undergoing protracted morphine withdrawal. Specifically, we sought to determine whether a prior history of chronic morphine exposure, compared to opioid-naïve mice, would induce enduring alterations in patterns of neural activation in response to a rewarding dose of a morphine challenge injection (Do Couto et al., 2003). To this end, within the same behavioral cohort from Figure 1, we examined FOS expression across 22 regions that are structurally connected to VTA^MOR^ neurons or involved in withdrawal processes following a morphine challenge administered 90 minutes prior to tissue collection **(Fig. 4a, Extended Data Table 4-1)**.

**Figure 4:**
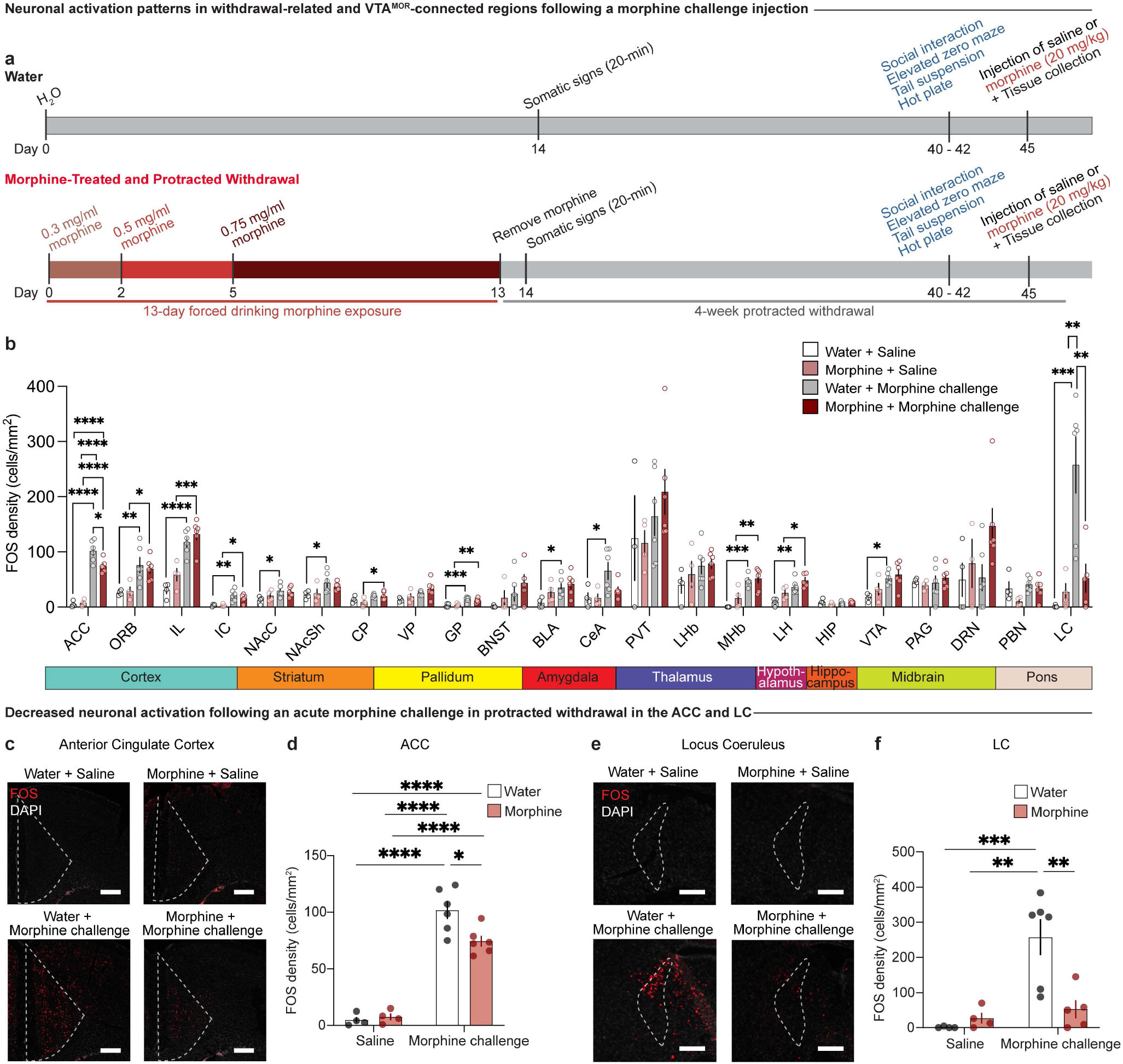
Protracted morphine withdrawal leads to decreased neuronal activation following acute morphine challenge in the ACC and LC. **(a)** Experimental schematic of the 13-day chronic forced morphine drinking exposure with escalating concentrations, assessment of somatic withdrawal signs 24-hours following cessation of morphine administration, a 4-week protracted withdrawal period, behavioral battery assessment, and tissue collection 90 minutes following a morphine challenge injection. **(b)** Neuronal activation as represented by FOS density in opioid naïve and morphine-treated mice following a saline or morphine challenge injection (20 mg/kg, s.c.) across 22 regions connected to VTA^MOR^ neurons or associated with withdrawal. Chronic morphine exposure results in decreased neuronal activation following an acute morphine challenge in protracted withdrawal in the **(c and d)** anterior cingulate cortex (two-way ANOVA + Bonferroni, treatment F_1,16_=4.312, p=0.0543; injection F_1,16_=190.3, p<0.0001; interaction F_1,16_=6.456, p=0.0218, scale, 200 μm) and **(e and f)** locus coeruleus (two-way ANOVA + Bonferroni’s multiple comparison, treatment F_1,15_=5.847, p=0.0288; injection F_1,15_=14.48, p=0.0017; interaction F_1,15_=9.757, p=0.0070, scale, 200 μm). Data are represented as mean ± SEM, *p<0.05, **p<0.01, ***p<0.001, ****p<0.0001.

Opioid-naïve mice displayed increased FOS expression in numerous regions following an acute morphine injection relative to saline-injected mice, including the ORB, ACC, IL, IC, NAcC, NAcSh, GP, CeA, BLA, MHb, LH, VTA, and LC **(Fig. 4b)**. This widespread activation is consistent with our lab’s previous findings that acute morphine elicits robust FOS expression in opioid naïve mice (Brynildsen et al., 2020). Our finding that an acute morphine injection increases FOS in the VTA of opioid naïve animals **(Fig. 4b)** may be largely attributed to the activation of dopaminergic neurons within this region (Georges et al., 2006; Corre et al., 2018). The VTA represents a key component of the mesocorticolimbic dopamine circuitry, and prior studies have found that morphine increases the impulse activity and burst firing of VTA dopamine neurons (Georges et al., 2006) and that opioids such as heroin activate dopaminergic neurons in the medial VTA (Corre et al., 2018). In mice with a prior history of chronic morphine exposure, an acute morphine injection induced heightened FOS density relative to saline injected animals, albeit in a more focused set of regions including the ORB, ACC, IL, IC, CP, GP, MHb, and LH Interestingly, in mice with a prior history of chronic morphine exposure, a morphine challenge injection during protracted withdrawal induced blunted FOS expression relative to opioid naïve mice in the ACC **(Fig. 4c-d;** two-way ANOVA + Bonferroni’s multiple comparison, treatment F_1,16_=4.312, p=0.0543; injection F_1,16_=190.3, p<0.0001; interaction F_1,16_=6.456, p=0.0218) and in the LC **(Fig. 4e-f;** two-way ANOVA + Bonferroni’s multiple comparison, treatment F_1,15_=5.847, p=0.0288; injection F_1,15_=14.48, p=0.0017; interaction F_1,15_=9.757, p=0.0070). This decreased neuronal activation in response to a rewarding dose of morphine (Do Couto et al., 2003) suggests persistent neuroadaptive changes resulting from chronic morphine exposure and subsequent 4-week withdrawal. These changes may contribute to dysregulated neural activity underlying maladaptive negative af-fect behaviors observed during withdrawal. These findings suggest that the ACC and LC may be uniquely susceptible to chronic morphine and protracted withdrawal-induced plasticity, potentially contributing to the persistence of negative affective states during withdrawal.

### VTA^MORs^ mediate low sociability during protracted morphine withdrawal

Our investigations have highlighted VTA^MOR^ neurons’ increased activation following the development of dependence and at the onset of withdrawal. Furthermore, our mapping of the baseline structural connectivity of VTA^MOR^ neurons to diverse projections targets implicated in reward, stress, and aversion indicates that this population could contribute to the negative affect behaviors characteristic of protracted withdrawal. Building on these findings, we next sought to directly assess the sufficiency of VTA^MORs^ in mediating protracted withdrawal-induced negative affect behaviors.

To re-express MORs specifically in the VTA, we performed bilateral injections of an AAV expressing a Cre-dependent human MOR (AAV-hMOR; AAVDJ-*hSyn1*-FLEX-mCh-T2A-FLAG-hMOR-WPRE) or a control virus (AAV-EGFP; AAV5-*hSyn*-FLEX-EGFP) into the VTA of Cre-expressing MOR KO mice **(Fig. 5a, b)**. To validate our viral VTA^MOR^ re-expression in the MOR KO line, we used quantitative PCR (qPCR) and confirmed reduced expression of mouse *Oprm1* in this line **(Fig. 5c**, unpaired t test, t=8.949, df=10, p<0.0001). Further qPCR analysis confirmed the viral-mediated re-expression of human *OPRM1* in the VTA of MOR KO mice **(Fig. 5d**, unpaired t test, t=8.551, df=5, p=0.0004), shown through increased human *OPRM1* expression compared to baseline levels in MOR KO mice.

**Figure 5:**
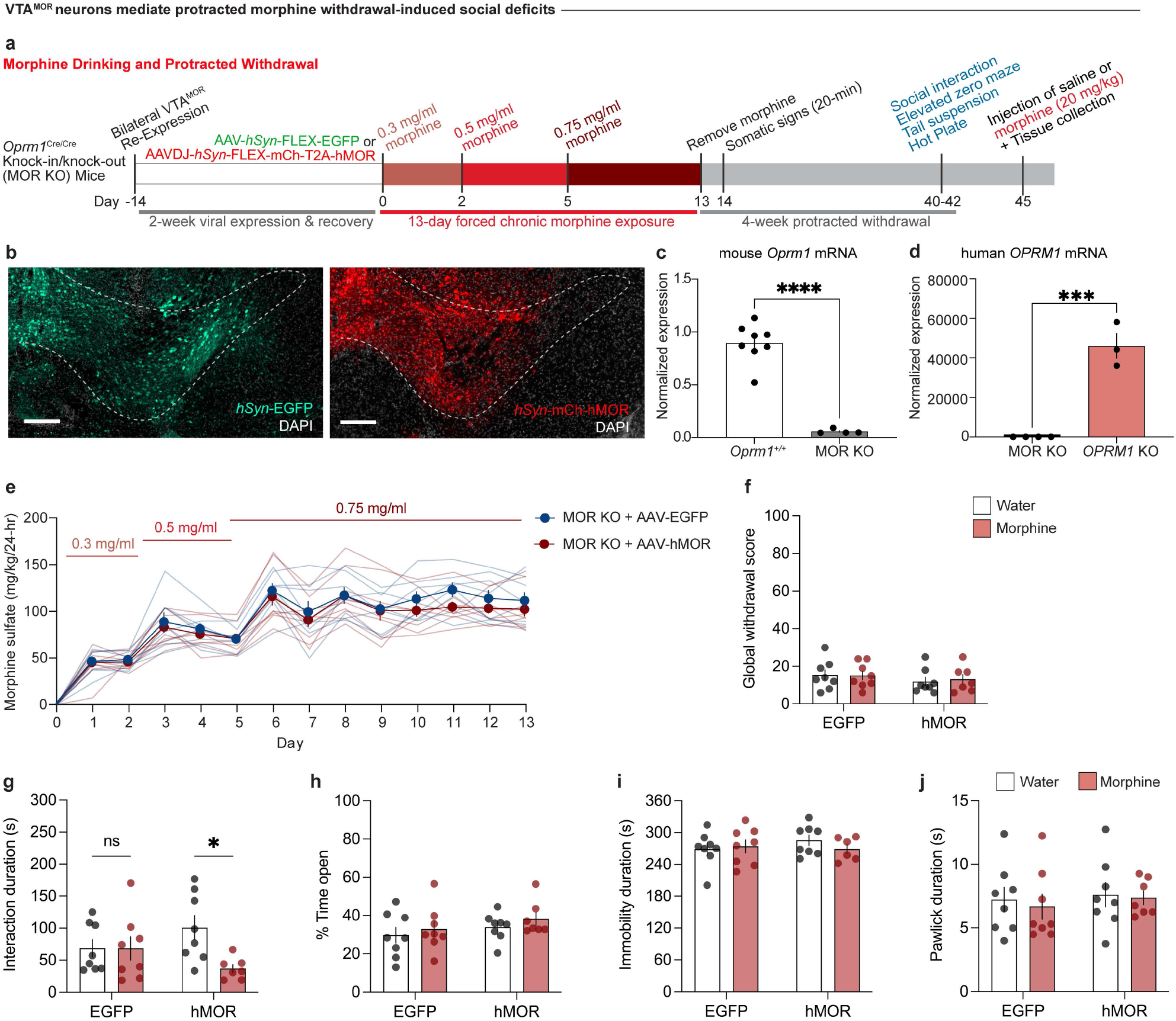
VTA^MORs^ mediate protracted withdrawal-induced low sociability. **(a)** Experimental schematic for VTA^MOR^ re-expression in MOR KO mice through bilateral VTA injections of AAVDJ-*hSyn1*-FLEX-mCh-T2A-FLAG-hMOR, followed by the protracted morphine withdrawal paradigm and culminating in a morphine challenge injection (20 mg/kg, s.c.). **(b)** AAV-*hSyn*-EGFP viral control or AAVDJ-*hSyn1*-hMOR in the VTA injection site (scale, 200 μm). **(c)** MOR knockout in the MOR KO mice demonstrated by decreased mouse *Oprm1* mRNA expression relative to the line’s littermate controls (n=8 *Oprm1*^+/+^ mice/4 MOR KO mice, unpaired t test, t=8.949, df=10, p<0.0001). **(d)** VTA^MOR^ re-expression in MOR KO mice through increased expression of human *OPRM1* mRNA relative to MOR KOs (n=4 MOR KO mice/3 MOR KO mice with VTA^MOR^ bilateral re-expression, unpaired t test, t=8.551, df=5, p=0.0004). **(e)** Daily morphine sulfate intake across a 13-day forced drinking exposure paradigm. **(f)** At 24 hours following morphine cessation, neither the AAV-EGFP-injected control or AAV-hMOR-injected treatment group show alterations in physical somatic signs as assessed through global withdrawal scores (n=8 AAV-EGFP-injected + opioid naïve mice/8 AAV-hMOR-injected + opioid naïve mice/8 AAV-EGFP-injected + morphine-treated mice/7 AAV-hMOR-injected + morphine-treated mice, two-way ANOVA + Bonferroni, virus F_1,27_=1.207, p=0.2817; treatment F_1,27_=0.04160, p=0.8399; interaction F_1,27_=0.09251, p=0.7633). **(g)** Morphine-treated MOR KO mice with VTA^MOR^ re-expression show decreased sociability relative to their opioid naïve counterparts during the 4-week protracted withdrawal period, while the morphine-treated AAV-EGFP-injected viral controls remain unaffected during protracted morphine (two-way ANOVA + Bonferroni, virus F_1,27_=0.0007256, p=0.9787; treatment F_1,27_=4.325, p=0.0472; interaction F_1,27_=4.250, p=0.0490). VTA^MOR^ re-expression in MOR KO mice does not alter protracted morphine withdrawal behaviors related to **(h)** Elevated zero maze (two-way ANOVA + Bonferroni, virus F_1,27_=1.727, p=0.1999; treatment F_1,27_=1.075, p=0.3090; interaction F_1,27_=0.01890, p=0.8917), **(i)** Tail suspension (two-way ANOVA + Bonferroni, virus F_1,26_=0.2659, p=0.6105; treatment F_1,26_=0.2668, p=0.6098; interaction F_1,26_=0.9168, p=0.3471), and **(j)** Nociceptive hot plate test (two-way ANOVA + Bonferroni, virus F_1,27_=0.3573, p=0.5550; treatment F_1,27_=0.1854, p=0.6702; interaction F_1,27_=0.03435, p=0.8544). Data = mean ± SEM, *p<0.05, ***p<0.001, ****p<0.0001.

Mice were exposed to the 13-day forced morphine drinking paradigm with escalating concentrations **(Fig. 5a, e)**. Morphine-treated mice in both the AAV-hMOR and AAV-EGFP injected groups did not show an increased consumption of morphine solution throughout the exposure paradigm relative to day 1 **(Extended Data Fig. 5a)**. This result contrasts with the morphine-treated wild-type C57BL/6J mice, which have functional MORs globally and exhibit a significant increase in morphine solution consumption relative to day 1 **(Fig. 1b)**. We assessed somatic withdrawal signs 24 hours following the cessation of morphine administration and found that neither the AAV-EGFP nor the AAV-hMOR injected groups showed differences in global withdrawal score between water and morphine-treated animals **(Fig. 5f;** two-way ANOVA + Bonferroni’s multiple comparison, virus F_1,27_=1.207, p=0.2817; treatment F_1,27_=0.04160, p=0.8399; interaction F_1,27_=0.09251, p=0.7633). While the VTA^MOR^ population is a critical pharmacological target of morphine, this result suggests that MORs in the VTA are not sufficient to induce spontaneous morphine withdrawal-induced physical dependence signs.

At 4 weeks of protracted morphine withdrawal, morphine-treated mice with VTA-specific re-expression of AAV-hMOR demonstrated decreased sociability relative to their opioid-naïve counterparts, while the AAV-EGFP-injected group, which effectively lack functional MORs, remain unaffected compared to opioid naïve mice **(Fig. 5g;** two-way ANOVA + Bonferroni’s multiple comparison, virus F_1,27_=0.0007256, p=0.9787; treatment F_1,27_=4.325, p=0.0472; interaction F_1,27_=4.250, p=0.0490). This outcome highlights the sufficiency of VTA^MORs^ in mediating low sociability during protracted morphine withdrawal. Additional behavioral assessments including the elevated zero maze **(Fig. 5h;** two-way ANOVA + Bonferroni’s multiple comparison, virus F_1,27_=1.727, p=0.1999; treatment F_1,27_=1.075, p=0.3090; interaction F_1,27_=0.01890, p=0.8917), tail suspension test **(Fig. 5i;** two-way ANOVA + Bonferroni’s multiple comparison, virus F_1,26_=0.2659, p=0.6105; treatment F_1,26_=0.2668, p=0.6098; interaction F_1,26_=0.9168, p=0.3471), and hot plate test **(Fig. 5j;** two-way ANOVA + Bonferroni’s multiple comparison, virus F_1,27_=0.3573, p=0.5550; treatment F_1,27_=0.1854, p=0.6702; interaction F_1,27_=0.03435, p=0.8544) did not reveal differences between morphine-treated and opioid-naïve mice within the AAV-EGFP and AAV-hMOR injected groups. These findings suggest that the effects of chronic morphine exposure on VTA^MORs^ may primarily influence sociability.

### VTA^MORs^ contribute to acute morphine-induced neural activation in the ACC

Given our previous findings of blunted FOS expression in the ACC and LC in morphine-treated C57 mice during protracted withdrawal, **(Fig. 4c-f)**, we sought to assess whether re-expression of VTA^MORs^ in MOR KO mice may modulate FOS expression in these regions following a morphine challenge. Interestingly, in opioid naïve MOR KO mice with VTA^MOR^ re-expression, FOS expression in the ACC is increased following an acute morphine challenge relative to the saline controls **(Fig. 6a, b;** three-way ANOVA + Bonferroni’s multiple comparison, treatment F_1,23_=0.5316, p=0.4733; virus F_1,23_=0.8245, p=0.3733; injection F_1,23_=7.059, p=0.0141; interaction (injection x virus x drinking F_1,23_=8.982, p=0.0064). This effect contrasts with results in opioid naïve group injected with AAV-EGFP, where FOS expression remains unaltered following the morphine challenge injection relative to the saline controls. This finding highlights the role of VTA^MORs^ in modulating the ACC’s neural activation response to a reinforcing dose of a morphine stimulus (Do Couto et al., 2003). Furthermore, in mice undergoing 4-week protracted morphine withdrawal with VTA^MOR^ re-expression, the acute morphine challenge induces decreased FOS relative to their opioid-naïve counterparts **(Fig. 6a, b)**. This effect suggests that chronic activation of VTA^MORs^ during the chronic morphine exposure, alongside the subsequent 4-week withdrawal, alters the ACC’s neural responsiveness to morphine.

**Figure 6:**
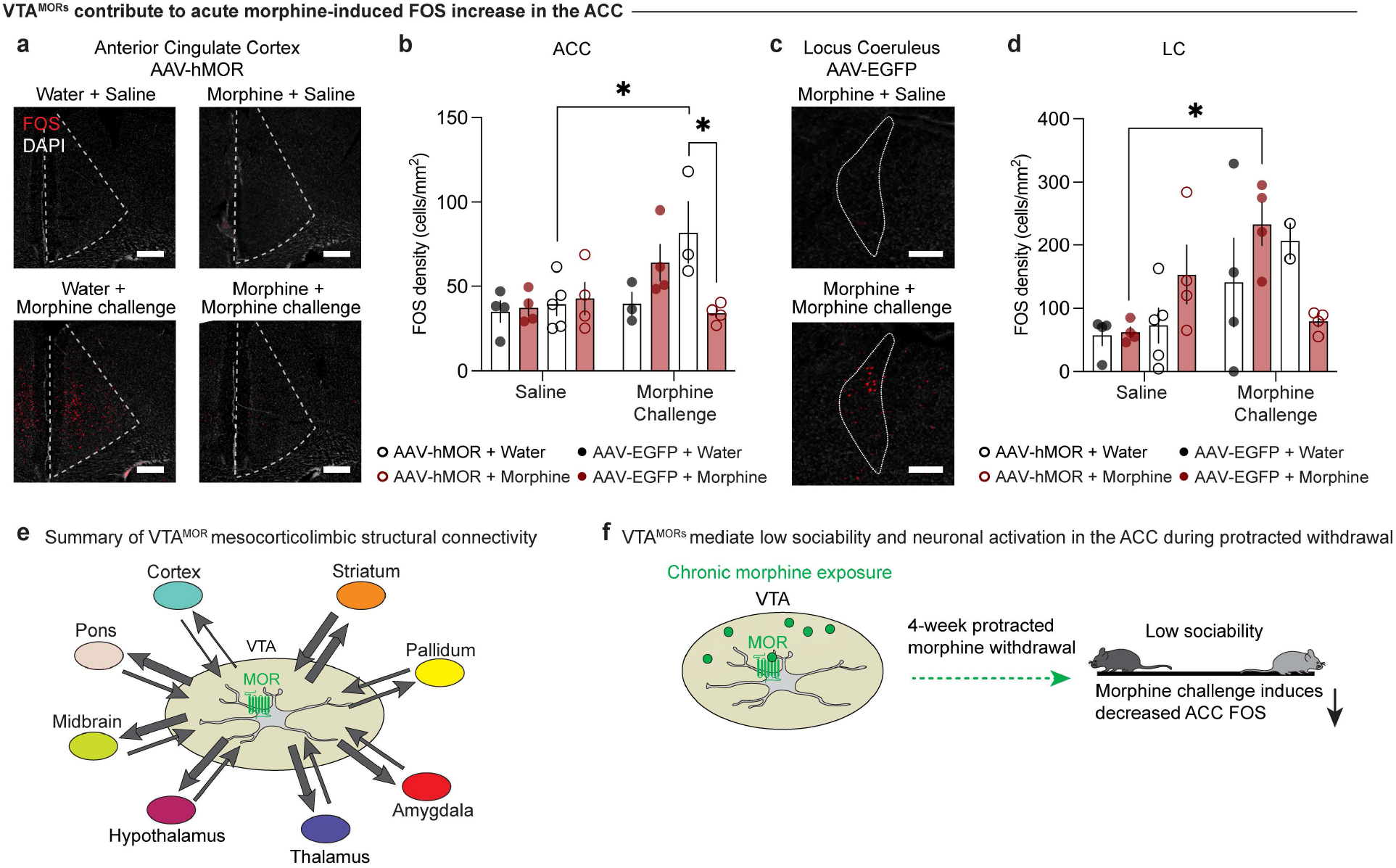
VTA^MORs^ contribute to acute morphine-induced FOS increase in the ACC. FOS density following a saline or morphine challenge injection (20 mg/kg, s.c.) in the **(a, b)** anterior cingulate cortex (ACC) (three-way ANOVA + Bonferroni’s multiple comparison, treatment F_1,23_=0.5316, p=0.4733; virus F_1,23_=0.8245, p=0.3733; injection F_1,23_=7.059, p=0.0141; interaction (injection x virus x drinking F_1,23_=8.982, p=0.0064, scale, 200 μm) and **(c, d)** locus coeruleus (LC) (three-way ANOVA + Bonferroni’s multiple comparison, treatment F_1,23_=0.2204, p=0.6431; virus F_1,23_=0.02855, p=0.8673; injection F_1,23_=8.597, p=0.0075; interaction (injection x virus x drinking F_1,23_=7.507, p=0.0117, scale, 200 μm). In the ACC, a morphine challenge injection leads to increased **(a, b)** FOS in AAV-hMOR-injected opioid naïve mice, while morphine-treated mice show reduced FOS expression compared to their opioid naïve counterparts. In the LC, a morphine challenge injection leads to increased **(c, d)** FOS in AAV-EGFP-injected opioid naïve mice. **(e)** Schematic representation of VTA^MOR^ neurons’ structural connectivity as explored across our VTA^MOR^ input-output mapping experiments. Arrow thickness represents estimated connection density. **(f)** Schematic summary of morphine bound to a VTA^MOR^ neuron and its subsequent effects on low sociability and morphine challenge-induced ACC FOS during protracted withdrawal. Data are represented as mean ± SEM, *p<0.05.

Additionally, we observed altered neuronal activation patterns in the LC in morphine-treated MOR KO mice injected with the AAV-EGFP virus. Following an acute morphine challenge injection, these MOR KO mice exhibited increased FOS expression in the LC **(Fig. 6c, d;** three-way ANOVA + Bonferroni’s multiple comparison, treatment F_1,23_=0.2204, p=0.6431; virus F_1,23_=0.02855, p=0.8673; injection F_1,23_=8.597, p=0.0075; interaction (injection x virus x drinking F_1,23_=7.507, p=0.0117). Given this effect was not observed in morphine-treated C57BL/6J mice following a morphine challenge injection (**Fig. 4e-f**), this result may suggest compensatory changes in response to morphine that may be mediated by alterations in noradrenergic signaling, as the locus coeruleus is the major source of norepinephrine (Berridge and Waterhouse, 2003). The differential response in morphine-treated mice that is not apparent in the opioid naive mice may also suggest unique opioid-induced plasticity in this area following chronic morphine exposure and protracted withdrawal.

## DISCUSSION

In this study, we aimed to elucidate the role of MORs in the VTA in facilitating neuroadaptive changes and negative affective behaviors during protracted morphine withdrawal. We first modeled the emergence of negative affective behaviors through increased social deficit-, anxiety-, and despair-related behaviors during protracted withdrawal and identified VTA^MOR^ neurons’ involvement in the onset of withdrawal across the VTA’s anterior to posterior axis. These VTA^MOR^ neurons are structurally connected to an array of cortical, subcortical and hindbrain regions central to motivation, withdrawal, affect, and reward, suggesting that withdrawal-related maladaptations in VTA^MOR^ neurons might affect and impact reward processes in upstream and downstream circuits. Indeed, chronic morphine exposure diminishes the ability of an acute, rewarding-dose of morphine to engage these regions during the protracted withdrawal period, in particular the ACC and LC. We also demonstrate that VTA^MORs^ are sufficient to facilitate low sociability during protracted withdrawal and contribute to the diminished neural activity in the ACC. Thus, our findings offer a neuronal-type, region-specific target implicated in a maladaptive phenotype of protracted opioid withdrawal – low sociability and diminished opioid-reward responsiveness (Chou et al., 2011).

Consistent with prior literature, the observed negative affective behaviors following four weeks of protracted morphine withdrawal parallel findings from various opioid administration paradigms. For instance, protracted withdrawal three to six weeks following investigator-administered opioid injections also induces low sociability (Goeldner et al., 2011; Bravo et al., 2020; Becker et al., 2021; Pomrenze et al., 2022), despair-like behaviors (Goeldner et al., 2011; Becker et al., 2021), and anxiety-related behaviors (Bravo et al., 2020; Becker et al., 2021), which reinforces the robust nature of these maladaptive behaviors as key features of protracted opioid withdrawal’s pathology. Our adoption of an oral morphine drinking paradigm in the home cage minimizes potential restraint stress during frequent injections (Stuart and Robinson, 2015) and mirrors the pharmacokinetic properties of oral opioid consumption in humans that enable a naturalistic framework of morphine dose escalation and maintenance.

Furthermore, VTA^MOR^ neurons do not appear to mediate somatic signs during protracted morphine withdrawal. Our finding that both morphine-treated AAV-hMOR and AAV-EGFP injected groups exhibited no differences in physical withdrawal signs relative to the opioid naïve mice suggests that VTA^MORs^ do not mediate physical signs of morphine withdrawal. This finding aligns with established literature that MORs in other regions, including the locus coeruleus (Maldonado et al., 1992; Maldonado and Koob, 1993) and central amygdala (Chaudun et al., 2024), are critical in mediating the physical signs of opioid withdrawal. Additionally, studies have shown that knockdown of MORs in the VTA in floxed *Oprm1*^*fl/fl*^ mice, which possess MORs flanked by *loxP* sites, does not alter the physical somatic signs characteristic of fentanyl withdrawal. This finding further supports our observation of no differences in physical withdrawal signs following morphine cessation.

Our findings reveal the nuanced contribution of VTA^MOR^ neurons in social interaction during protracted withdrawal. This result extends beyond the established influence of dopaminergic transmission (Solié et al., 2022) and oxytocin signaling (Hung et al., 2017) in social reward in the VTA. Dysregulation of MOR activity is implicated in a variety of neuropsychiatric disorders, such as depression, anxiety and substance use disorders, where social behavior is often adversely affected (Fone and Porkess, 2008; Nummenmaa et al., 2020; Christie, 2021). The reduced sociability previously observed in other models of MOR KO mice underscores MORs’ integral function in modulating social behavior (Moles et al., 2004; Becker et al., 2021; Toddes et al., 2021), a role that may be disrupted during repeated opioid exposure and protracted abstinence. The involvement of MOR-expressing neurons in negative affect has been investigated within regions such as the dorsal raphe nucleus (DRN) (Lutz et al., 2014; Bailly et al., 2023) and habenula (Bailly et al., 2023), as the deletion of MORs in the DRN prevents the emergence of social deficits during chronic heroin withdrawal (Lutz et al., 2014). Moreover, activation of MOR neurons in the habenula mediates projection-specific aversion, as optogenetic stimulation of habenula MOR neurons projecting to the DRN increase anxiety-like behavior, while optogenetic stimulation of these neurons projecting to the interpeduncular nucleus (IPN) produces despair-related behavior and avoidance (Bailly et al., 2023). Despite the established role of the VTA as a key target for opioids, the field’s understanding of the behavioral implications of VTA^MOR^ neurons during dependence and protracted withdrawal remains incomplete. Our work addresses this gap by demonstrating this neuronal population’s role in facilitating social deficit-related behavior during withdrawal. Interestingly, other behavioral measures such as anxiety-, despair, and nociceptive-related behaviors did not show differences between morphine-treated and opioid naïve mice upon VTA^MOR^ viral re-expression, suggesting that MORs in the VTA may be more specific to social behaviors during protracted withdrawal.

While we have identified the role of VTA^MORs^ in facilitating low sociability during protracted morphine withdrawal, an important consideration to acknowledge is the use of certain strains of MOR KO mice, which have been previously reported to exhibit low sociability (Moles et al., 2004; Becker et al., 2014; Toddes et al., 2021; Derieux et al., 2022). However, it is worth noting that the MOR KO line used in these previous studies (Matthes et al., 1996) was generated by Dr. Brigitte Kieffer using different technologies and knock-in strategies (129S2/SvPas x C57BL/6-derived P1 ES cells inserted into exon 2), compared to the MOR KO line used here, which was created by Dr. Richard Palmiter (129S6 x C57BL/7-derived F1-derived G4 ES cells through insertion of the Cre:GFP construct into exon 1). To our knowledge, the MOR KO line generated by Dr. Richard Palmiter’s lab, used in this study, has not been reported to exhibit such social deficits, nor did we observe any social behavior deficits in this line (**Extended Data 5-1l;** unpaired t test, t=0.6239, df=19, p=0.5401). Differences in genetic background or environmental factors, such as vivarium conditions or experimental procedures, could contribute to behavioral discrepancies and variability in phenotypic outcomes (Crabbe et al., 1999). Moreover, our primary objective was to investigate the role VTA-specific MORs in mediating social behaviors following chronic opioid exposure and withdrawal. The use of Cre-dependent viral tools enabled the selective re-expression of MORs within the VTA of MOR KO mice to dissect the specific contributions of this neural population to withdrawal-related negative affective behaviors. Future studies to further elucidate the contribution of VTA^MORs^ in facilitating withdrawal-related social behaviors include utilizing conditional knockout models and knocking down MORs within the VTA of floxed Oprm1^fl/fl^ mice. Additionally, expanding the behavioral repertoire of social behaviors through other assessments, such as the three-chamber social test or social conditioned place preference, could provide an enhanced understanding of the role of VTA^MORs^ in diverse social contexts.

We also found that mice with a prior history of morphine exposure exhibit decreased neuronal activation in the ACC following an acute morphine injection. The ACC has been implicated in encoding and multiplexing various aspects of reward processing (Hayden and Platt, 2010; Cai and Padoa-Schioppa, 2012; Monosov, 2017), including social interaction (Rudebeck et al., 2008). In our study, we observe blunted morphine challenge-induced ACC FOS in mice undergoing protracted morphine withdrawal. The diminished neural responsiveness in the ACC to a reinforcing dose of morphine (Do Couto et al., 2003) may reflect a dysregulation in processing rewarding stimuli including social reward, an effect that may inform our observation of social deficits during protracted withdrawal. Specifically, our results shed light on the role of VTA^MOR^ neurons in modulating the ACC’s response to acute morphine **(Fig. 6a, b)**. Despite the relatively minimal direct structural connections between VTA^MOR^ neurons and the ACC, as observed in our rabies input mapping **(Fig. 3d)** and axonal projection data **(Fig. 3h)**, increased ACC FOS following a morphine challenge in MOR KO mice with restored VTA^MOR^ expression suggests that indirect or neuromodulatory pathways may mediate this effect. Opioid-induced MOR activation induces downstream effects, notably the inhibition of local GABAergic interneurons within the VTA (Johnson and North, 1992). Given the characterized projection of dopaminergic VTA neurons to the ACC (Beier et al., 2015; Breton et al., 2019), it is plausible that this altered dopaminergic signaling following acute morphine may contribute to the heightened FOS density in the ACC. This aligns with prior evidence that VTA dopaminergic projections to the ACC are critical for the acquisition and maintenance of morphine conditioned place preference (Narita et al., 2010). Moreover, the variety of neuronal types within the VTA, such as glutamate-transmitting neurons (Yamaguchi et al., 2007; Nair-Roberts et al., 2008; Root et al., 2020), may also contribute to morphine-induced FOS expression. However, the projection targets of VTA glutamatergic neurons remain to be elucidated.

In conclusion, our study highlights the role of VTA^MORs^ in mediating low sociability during protracted withdrawal. We demonstrate that chronic morphine exposure leads to increased negative affective behaviors during withdrawal, which are linked to heightened neuronal activity in VTA^MOR^ neurons and their structurally connected regions implicated in reward and aversion processing. By highlighting the region- and neuronal-specific mechanisms underlying neuroadaptive alterations from chronic opioid exposure and withdrawal, this study advances our understanding of neural substrates involved in dependence and withdrawal. Future research may expand investigations of VTA^MOR^ neurons’ role in protracted withdrawal-induced low sociability, as this neuronal population may represent a potential target for interventions aimed at alleviating the debilitating symptoms associated with protracted abstinence.

## ACKNOWLEDGEMENTS

This research was funded by NIH R01-DA-054374, R01-DA-056599, DP2-GM1-40923, and T32-DA028874. We thank the University Laboratory Animal Resources (ULAR) group at the University of Pennsylvania’s Translational Research Laboratory for assistance with rodent husbandry and veterinary support. We thank additional members of the Corder and Blendy Labs, including Blake Kimmey, Lisa Woolridge, and Malaika Mahmood for technical assistance. We also thank Matthew Banghart and Karl Deisseroth for their virus contributions.

## AUTHOR CONTRIBUTIONS

A.J., G.C., J.B. conceptualized and planned out the study. A.J. performed all stereotaxic surgeries, morphine treatment paradigms, behavioral experiments, tissue processing, immunohistochemistry, imaging, and wrote the original manuscript. Y.X. performed the cell quantification. R.A.S.O. developed the AxoDen algorithm for axonal density quantification. K.B. conceptualized and designed the rabies viral construct. A.R. and K.T.C. performed the qPCR. All authors contributed to the editing and revising of the document.

## COMPETING INTERESTS

The authors declare no competing financial interests.

## MATERIALS AND METHODS

### Animals

Male and female mice aged 8–20-week-old were used from the following genetic lines: C57BL/6J mice were purchased from Taconic Biosciences and bred in-house at the University of Pennsylvania. A MOR-specific T2A-cleaved Cre-recombinase (Mengaziol et al., 2022) mouse line that will be referred to as *Oprm1*^MOR-T2A-Cre^ (C57BL/6NTac-^Oprm1em1(cre)Jabl^/Mmnc; Stock #070963-UNC) was generated and bred-inhouse; briefly, the line was generated by inserting a T2A cleavable peptide sequence and the Cre coding sequence into the MOR 3’UTR, as previously described (Mengaziol et al., 2022). An *Oprm1*^Cre:GFP^ knock-in/knock-out mouse line that will be referred to as MOR KO (B6.Cg-*Oprm1*^*tm1*.*1(cre/GFP)Rpa*^/J, Stock # #035574) was generated from Dr. Richard Palmiter’s laboratory and bred in-house; briefly, the MOR KO line was generated by inserting a cassette encoding Cre:GFP 5’ of the initiation codon in *Oprm1*’s first coding exon, as previously described (Liu et al., 2021), and mice homozygous for Cre:GFP lack MOR expression and therefore do not respond to endogenous or exogenous ligands. For both the *Oprm1*^MOR-T2A-Cre^ and MOR KO lines, only mice homozygous for Cre were used for behavioral studies. Wild-type controls of the MOR KO line, *Oprm1*^+/+^, which lack the Cre allele, were used to validate expression using qPCR. Mice were maintained on a 12-hour reverse light/dark cycle (lights on at 6 pm) and provided with food and water *ad libitum*. All mice in the drinking experiments were singly-housed to measure individual daily volume consumption. All conducted experiments were approved by the Institutional Animal Care and Use Committee of the University of Pennsylvania.

### Stereotaxic Surgeries and Viral Injections

Mice were anesthetized with isoflurane (2.5-3.0% for induction, 1.5-2.0% for maintenance) and head-fixed in a stereotaxic frame. For brain-wide MOR+ neuron labelling, AAV.PHP.eB-*mMORp*-eYFP (titer: 8.60E+11 GC/mL) was administered intravenously through the retro-orbital sinus using a 30 G insulin syringe, as previously described (Salimando et al., 2023). Other viruses in the study were injected intracranially into the VTA (AP: −3.2 mm, ML: ±0.55 mm, DV: −4.75 mm) using a 10 μL Nanofil Hamilton syringe (WPI) with a 33 G beveled needle. For the rabies monosynaptic labelling, 150 nL of a 1:1 mixture of AAV5-*CAG*-FLEX-TVA-mCherry and AAV8-*CAG*-FLEX-RABV-G (titer: 1.00E+12 GC/mL, Beier Lab) was unilaterally injected into the left VTA of *Oprm1*^MOR-T2A-Cre^ mice. Two weeks post-injection, 300 nL of RABVΔG-GFP (titer: 8.00E+08 GC/mL, Beier Lab) was administered into the same injection site and mice were euthanized 5 days later. To map axonal density outputs of VTA^MOR^ neurons, 150 nL of AAV5-*CAG*-FLEX-tdTomato (titer: 2.10E+13 GC/mL, Addgene, cat. no. 28306-AAV5) was unilaterally injected into the left VTA of *Oprm1*^MOR-T2A-Cre^ mice. For re-expression of VTA^MORs^, MOR KO mice received bilateral VTA injections of 300 nL per side of AAVDJ-*hSyn1*-FLEX-mCh-T2A-FLAG-hMOR-WPRE (AAV-hMOR; titer: 1.30E+12 GC/mL, Banghart Lab) or control AAV5-*hSyn1*-DIO-EGFP (AAV-EGFP; titer: 1.30E+12 GC/mL, Addgene, cat. no. 50457-AAV5). Following surgery, mice received meloxicam (5 mg/kg, s.c.) and recovered for one to two weeks before behavioral assessments.

### Morphine Treatment

Morphine sulfate (NIDA Drug Supply, Research Triangle Park, NC) was dissolved in water at an initial concentration of 0.3 mg/mL for the first two days. The concentration of the morphine solution escalated to 0.5 mg/mL for the following three days and then maintained at 0.75 mg/mL for the remainder of the total 13-day exposure paradigm, as previously established and described (Belknap, 1990). We employed a forced drinking paradigm as the morphine solution was presented in 15-mL tubes within the home cage as the sole drinking source for continuous 24-hour access. Opioid naïve mice were administered water. Both the volume of the morphine solution and the mice’s body weights were recorded daily at 12:00 pm.

### Behavior

Mice were acclimated for 1 hour in the behavior room prior to all behavior assessments.

#### Somatic Withdrawal Signs

For precipitated withdrawal, mice were injected with naloxone (1 mg/kg, s.c.) on day 13 of the morphine exposure paradigm and subsequently placed in a clear observation box to monitor withdrawal signs. For spontaneous withdrawal, somatic signs were observed on day 14, 24 hours after cessation of morphine exposure. Somatic withdrawal signs were conducted for 20 minutes under indirect lighting (15 lux), during which the frequency of episodes of diarrhea, resting tremor, paw tremors, jumping, gnawing on limbs, genital licking and grooming, head shakes, body shakes, backing and defensive treading, and scratching were video-recorded and quantified, as previously described (Eacret et al., 2023) using an ethological keyboard in the BORIS (Friard and Gamba, 2016) software. The total frequency of each individual withdrawal behavior culminated to the global withdrawal score.

#### Social Interaction

Sex-, weight-, and age-matched unfamiliar mice that were drug-naïve were introduced into an open-field arena (40 x 40 cm) under indirect lighting (15 lux) for 10 minutes. Social interactions, including nose contacts (nose-to-nose, nose-to-body or nose-to-anogenital region), grooming, and following episodes, were scored using an ethological keyboard in the BORIS (Friard and Gamba, 2016) software, following established protocols (Goeldner et al., 2011; Lutz et al., 2014; Becker et al., 2021). The total duration of these episodes was summed as the social interaction duration.

#### Elevated Zero Maze

Mice were placed in an elevated zero maze (50 cm diameter, 5 cm track width, and 50 cm from the floor) under indirect lighting (15 lux) for 5 minutes. The total duration spent in the open and closed arms were recorded and analyzed using the Noldus Ethovision^®^ XT software (Leesburg, VA, USA).

#### Tail Suspension Test

Mice were suspended by the tail under indirect lighting (15 lux) for 6 minutes using a non-irritating adhesive tape. Total immobility duration was assessed, as previously described (Can et al., 2011).

#### Hot Plate Test

Mice were placed on a hot plate set to 50°C for 1 minute under indirect lighting (15 lux). The duration of hind paw and forepaw licking behaviors was recorded and analyzed using an ethological keyboard in BORIS (Friard and Gamba, 2016), and the total duration was recorded as the paw lick duration.

### Immunohistochemistry

90 minutes following an injection of saline, naloxone (1 mg/kg, s.c.; **Fig. 2a**), or an acute morphine challenge (20 mg/kg, s.c.; **Fig. 4a, 5a**), mice were anesthetized with sodium pentobarbital (50 mg/kg, i.p.) and perfused with ice-cold 0.01 M PBS followed by 4% paraformaldehyde (PFA). Brains were left in 4% PFA at 4°C overnight and subsequently transferred to 30% sucrose in PBS. Brains were then sectioned frozen at −20°C on a cryostat (Cryostar NX-50, ThermoScientific) and 30 μm thick coronal sections were collected and stored in 0.01 M PBS.

Sections were washed 3 × 10 minutes in 0.01 M PBS and then incubated for two hours in blocking buffer solution consisting of 5% normal donkey serum and 0.1% Triton X-100 in 0.01 M PBS. Sections were incubated overnight in room temperature with either rabbit anti-cFos (1:500, Cell Signaling Technologies, cat. no. 2250S), rabbit anti-dsRed (1:1000, Takara, cat. no. 632496), chicken anti-GFP (1:500, Abcam, cat. no. AB13970), or mouse anti-tyrosine hydroxylase (1:1000, Sigma-Aldrich, cat. no. MAB318) in the blocking buffer solution. Sections were then washed 3 x 10 minutes in 0.01 M PBS the following day and incubated with Alexa Fluor 647 donkey anti-rabbit (1:500, Invitrogen Thermo Fisher A31573), Alexa Fluor 594 donkey anti-rabbit (1:500, Invitrogen Thermo Fisher, cat. no. A21207), Alexa Fluor 488 donkey anti-chicken (1:500, Jackson Immuno Research, cat. no. 703-545-155), or Alexa Fluor 647 donkey anti-mouse (1:500, Invitrogen Thermo Fisher, cat. no. A31571) in the blocking buffer solution for 2 hours at room temperature. Sections were then washed 3 x 10 minutes in 0.01 M PBS, incubated in DAPI (1:10,000, Sigma-Aldrich, cat. no. D9542), mounted, and then cover slipped prior to imaging on a Keyence BZ-X810 fluorescence microscope.

### Quantification

#### Naloxone-induced FOS expression in VTA^MOR^ neurons

FOS- and YFP-labelled MOR neurons were quantified across the anterior to posterior axis of the VTA. 10 coronal VTA sections per mouse were assigned as anterior (−2.80 mm, −2.92 mm, and −3.08 mm from bregma), medial (−3.16 mm, −3.28 mm, and −3.4 mm from bregma), or posterior (−3.52 mm, −3.64 mm, −3.8 mm, and −3.88 mm from bregma) VTA sections, as based on the Franklin and Paxinos mouse brain atlas, 3^rd^ edition (Franklin and Paxinos, 2007). The percentage of FOS+ MOR+ neurons was calculated by dividing the total number of co-labelled FOS+ MOR+ neurons by the total number of MOR+ neurons in the VTA for each coronal section.

#### Acute morphine-induced FOS expression

FOS expression was analyzed in 22 regions implicated in withdrawal (Welsch et al., 2020) and identified as key VTA afferent and efferent projection targets (Beier et al., 2015). These regions included the orbital cortex (ORB), anterior cingulate cortex (ACC), infralimbic cortex (IL), insular cortex (IC), nucleus accumbens core (NAcCore), nucleus accumbens shell (NAcShell), caudate putamen (CP), globus palli-dus (GP), bed nucleus of the stria terminalis (BNST), ventral pallidum (VP), central amygdala (CeA), basolateral amygdala (BLA), hippocampus (HIP), paraventricular nucleus of the thalamus (PVT), lateral habenula (LHb), medial habenula (MHb), lateral hypothalamus (LH), ventral tegmental area (VTA), periaqueductal grey (PAG), dorsal raphe nucleus (DRN), parabrachial nucleus (PBN), and the locus coeruleus (LC). FOS expression was quantified and analyzed using a semi-manual detection method previously developed in our lab (Xie et al., 2024).

#### Rabies monosynaptic VTA^MOR^ input labelling

Every sixth 30-μm thick coronal section throughout the entire brain was included in analysis. GFP-positive VTA^MOR^ input cells from the left hemisphere were analyzed using Semi-Automated Workflow for Brain Slice Histology Alignment, Registration, and Cell Quantification (SHARCQ) (Lauridsen et al., 2022) and quantified, as previously described (Schwarz et al., 2015; Xie et al., 2024). Counts from each region were adjusted by a factor of 6 to account for the sampling rate, as previously reported (Beier et al., 2015). The fraction of GFP+ counts per region by the total sum of all rabies-labelled inputs throughout the entire brain was assessed for these regions chosen for analysis, which included major VTA inputs outside of the excluded region near the injection site, as previously described (Beier et al., 2015).

#### Axon density quantification of VTA^MOR^ outputs

Four 30-μm coronal sections per region were selected across the region’s anterior to posterior segment. An inclusive set of regions was selected based on their involvement in mediating opioid dependence and withdrawal (Welsch et al., 2020; Ozdemir et al., 2023), as well as their structural connectivity to the VTA (Beier et al., 2015). Coronal histology images were processed using AxoDen, a custom-written algorithm developed in the Corder Lab (https://corderlab.com) for the automated quantification of axonal density in defined regions (https://github.com/raqueladaia/AxoDen; web application link: (https://axoden.streamlit.app/) (Sandoval Ortega et al., 2024). Briefly, this algorithm converts images to gray scale and later employs dynamic thresholding to accurately binarize the image and segregate signal from background fluorescence. This method facilitates the quantification of the density of axon projections in selected brain regions.

### qPCR for mouse *Oprm1* and human *OPRM1*

Tissue samples from the VTA were collected as one mm punches from *Oprm1*^+/+^ mice (wild-type controls of the MOR KO line), MOR KO mice, and MOR KO mice with VTA^MOR^ re-expression through bilateral injections of AAVDJ-*hSyn1*-FLEX-mCh-T2A-FLAG-hMOR-WPRE (AAV-hMOR). Both the mouse *Oprm1* and human *OPRM1* transcripts were quantified. Total tissue RNA was extracted with RNAzol (Sigma-Aldrich, R4533) according to manufacturer’s protocol, and cDNA was synthesized from 1 ug RNA (Applied Biosystems, cat. no. 4374966). cDNA was diluted 1:10 and assessed for mRNA transcript levels by qPCR with SYBR Green Mix (Applied Biosystems, cat. no. A25741) on a QuantStudio7 Flex Real-Time PCR System (Thermo Fisher). Oligonucleotide primer sequences for target and reference genes are as follows: mouse_*Oprm1* (forward: CTGCAAGAGTTGCATGGACAG, reverse: TCAGATGACATTCACCTGCCAA); human_*OPRM1* (forward: ACTGATCGACTTGTCCCACTTAGATGGC, reverse: ACTGACTGACTGACCATGGGTCGGACAGGT); mouse_*L30* (forward: ATGGTGGCCGCAAAGAAGACGAA, reverse: CCTCAAAGCTGGACAGTTGTTGGCA). Target gene expression was normalized to an internal control (m*L30* mRNA) and sample control (*Oprm1*^*+/+*^) and analyzed by the 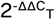 method (Livak and Schmittgen, 2001). Each sample reaction was performed in triplicate.

### Experimental Design and Statistical Analysis

#### Behavioral Assessments

Behavioral experiments were conducted to assess the effects of chronic morphine exposure on somatic withdrawal signs alongside negative affective behaviors. Mice underwent a 13-day morphine drinking paradigm followed by behavioral experiments including somatic withdrawal signs, social interaction, elevated zero maze, tail suspension test, and hot plate test. For C57BL/6J mice, 37 mice (n=23 males, n=14 females) were divided into two groups: opioid naive (n=16) and morphine-treated (n=21). Behavioral data were analyzed between opioid naïve and morphine-treated groups using unpaired t tests **(Fig. 1d-h and Fig. 2b)**. For MOR KO mice, 31 mice (n=22 males, n=9 females) were divided into four groups (n=7-8/group): AAV-EGFP virus + opioid naïve, AAV-EGFP virus + morphine-treated, AAV-hMOR virus + opioid naïve, and AAV-hMOR virus + morphine-treated **(Fig. 5f-j)**. A two-way ANOVA and Bonferroni’s multiple comparisons test was performed to analyze the factors of viral injection (AAV-hMOR vs. AAV-EGFP) and treatment condition (opioid naïve vs. morphine-treated). We acknowledge the sex imbalance in the MOR KO mice used in behavioral assessments due to constraints in the available MOR KO litters from our breeding colony.

#### Neuronal Activation Studies

Neuronal activation was assessed through FOS expression in male and female C57BL/6J and MOR KO mice at the culmination of behavioral assessments. To assess precipitated withdrawal-induced FOS expression in VTA^MOR^ neurons, mice were divided into two groups (n=3-5/group): opioid naïve + naloxone injection and morphine-treated + naloxone injection **(Fig. 2a)**. To assess potential subregional differences in neuronal activation within the VTA, 10 sections per mouse spanning the anterior to posterior VTA axis from −2.80 to −3.88 mm from bregma were analyzed. An unpaired t test was used to assess neuronal activation in the opioid naïve vs. morphine-treated groups **(Fig. 2d, g)** or two-way ANOVA and Bonferroni’s multiple comparison was performed to evaluate neuronal activation across treatment groups (opioid naïve vs. morphine-treated) and VTA subregions (anterior, medial, and posterior) **(Fig. 2e, h)** or specific bregma levels **(Fig. 2f, i)**. To assess the effects of a morphine challenge injection during protracted morphine withdrawal on neuronal activation, we analyzed FOS expression in 22 regions **(Fig. 4b)** after a morphine challenge during 4-week protracted withdrawal across four groups (n=4-6/group): opioid naïve + saline injection, morphine-treated + saline injection, opioid naïve + morphine challenge injection, and morphine-treated + morphine challenge injection. Two-way ANOVA and Bonferroni’s multiple comparisons was performed to evaluate factors of treatment (opioid naïve vs. morphine-treated) and injection (saline vs. morphine challenge) on neuronal activation **(Fig. 4 b-f)**. In MOR KO mice, neuronal activation in the ACC and LC was analyzed following a morphine challenge injection during the 4-week protracted withdrawal across 8 groups (n=3-5/group): AAV-EGFP + opioid naïve + saline injection, AAV-EGFP + morphine-treated + saline injection, AAV-hMOR + opioid naïve + saline injection, AAV-hMOR + morphine-treated + saline injection, AAV-EGFP + opioid naïve + morphine challenge injection, AAV-EGFP + morphine-treated + morphine challenge injection, AAV-hMOR + opioid naïve + morphine challenge injection, and AAV-hMOR + morphine-treated + morphine challenge injection **(Fig. 6 a-d)**. We performed a threeway ANOVA and Bonferroni’s multiple comparisons to assess viral groups (AAV-hMOR vs. AAV-EGFP), treatment conditions (opioid naïve vs. morphine-treated), and injection (saline vs. morphine challenge) on neuronal activation. We acknowledge the potential limitations in statistical power due to smaller sample sizes in groups, which were constrained by technical limitations of custom virus availability and breeding litters.

#### Structural Input-Output Connectivity Studies

For the VTA^MOR^ structural connectivity mapping, 6 *Oprm1*^MOR-T2A-Cre^ mice (n=4 males, n=2 females) were used for the rabies virus monosynaptic labelling (n=3) or axonal output mapping (n=3). Although the use of n=3 per group in our baseline connectivity mapping of VTA^MOR^ neurons is a lower sample size, this sample size is within range of established literature using similar tracing techniques (Beier et al., 2015, 2017) and our objective was to comprehensively map out the native structural connectivity of VTA^MOR^ neurons in an opioid-naïve state.

All data are presented as mean ± the standard error of the mean (SEM) for each group and analyzed using an unpaired t test or one-way, two-way, or three-way ANOVA and Bonferroni’s multiple comparison. Statistical analyses were performed using Prism 10 (GraphPad). Statistical outliers were identified using Grubbs’ test with an alpha = 0.05 significance level. One statistically significant outlier was excluded from the medial habenula FOS expression analysis in Figure 4, and one statistically significant outlier from the social interaction test was excluded from behavioral assessment in Figure 5.

## EXTENDED DATA

**Extended Data Figure 1-1:**
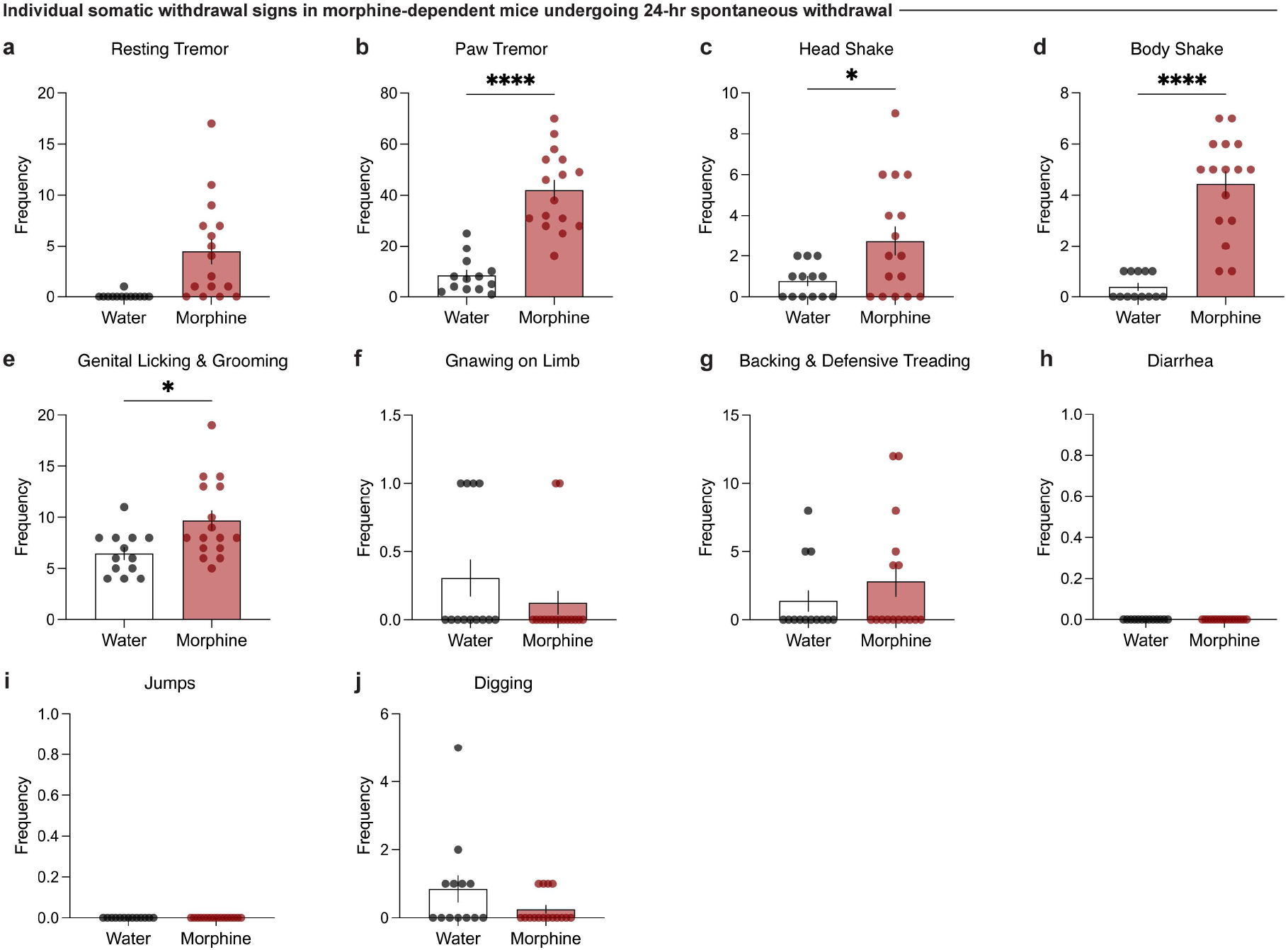
Individual somatic withdrawal signs in morphine-dependent mice undergoing 24-hour spontaneous withdrawal. **(a)** Morphine-dependent mice undergoing spontaneous withdrawal demonstrate increased frequencies of **(b)** paw tremors (unpaired t test, t=7.187, df=27, p<0.0001), **(c)** head shakes (unpaired t test, t=2.443, df=27, p=0.0214), and **(d)** body shakes (unpaired t test, t=7.342, df=27, p<0.0001), while the frequencies of **(a)** resting tremors, **(e)** genital licking and grooming, **(f)** gnawing on limb, and **(g)** backing and defensive treading bouts **(h)** diarrhea, **(i)** jumps, and **(j)** digging bouts remain unchanged relative to opioid naïve. Data are represented as mean ± SEM, *p<0.05, **p<0.01, ***p<0.001, ****p<0.0001.

**Extended Data Figure 2-1:**
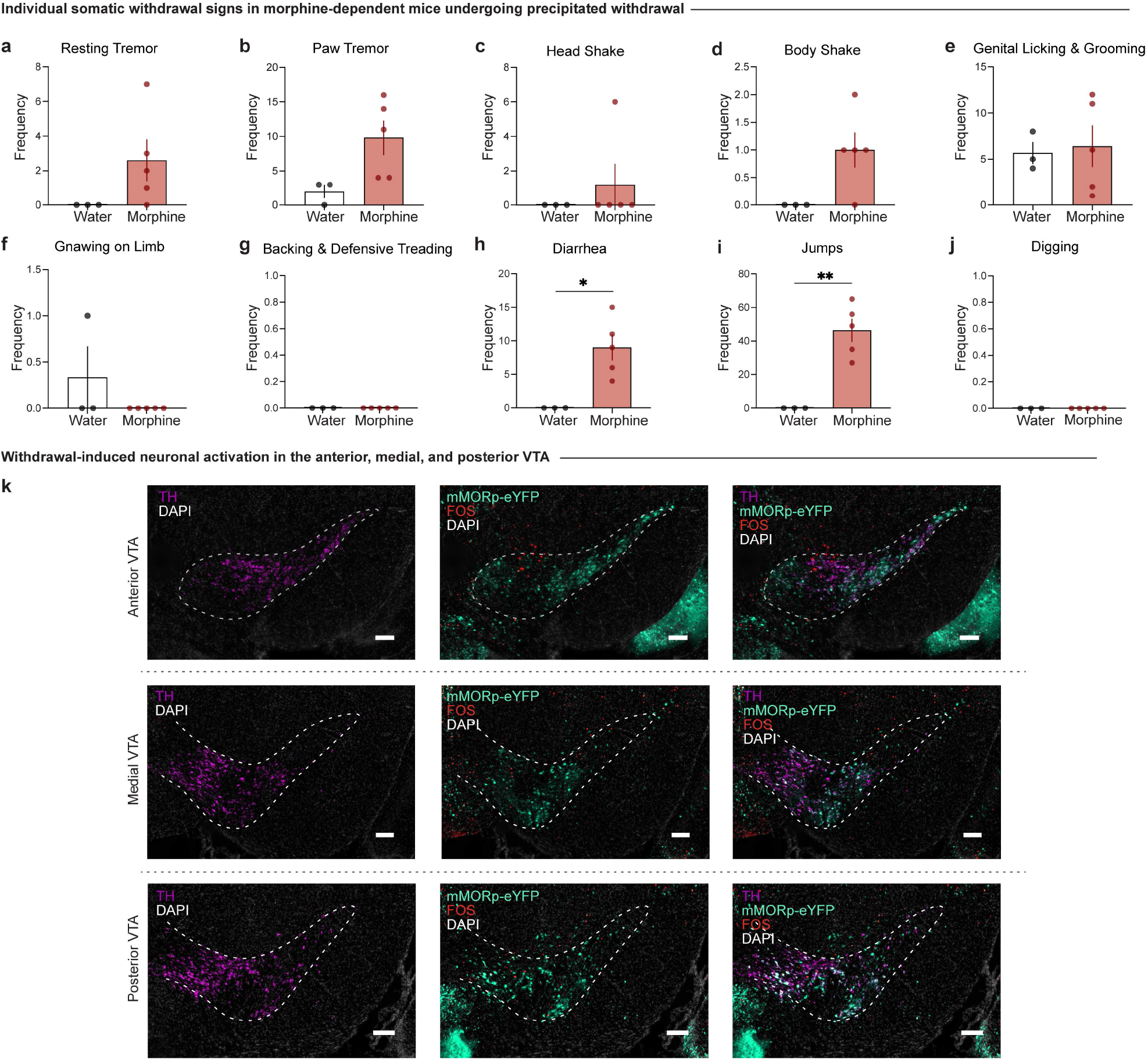
Individual somatic withdrawal signs in morphine-dependent mice undergoing precipitated withdrawal and neuronal activation of MOR neurons in the anterior, medial, and posterior VTA. Morphine-dependent mice undergoing precipitated withdrawal demonstrate increased frequencies of **(h)** diarrhea (unpaired t test, t=3.509, df=6, p=0.0127) and **(i)** jumps (unpaired t test, t=5.046, df=6, p=0.0023), while the frequencies of **(a)** resting tremors, **(b)** paw tremors, **(c)** head shakes, **(d)** body shakes, **(e)** genital licking and grooming, **(f)** gnawing on limb, and **(g)** backing and defensive treading bouts remained unchanged relative to opioid naïve mice. **(k)** Representative images of coronal sections showing tyrosine hydroxylase (TH) staining with *mMORp*-eYFP and FOS staining in the anterior, medial, and posterior VTA (scale, 200 μm). Data are represented as mean ± SEM, *p<0.05, **p<0.01. Scale, 200 μm.

**Extended Data Figure 3-1:**
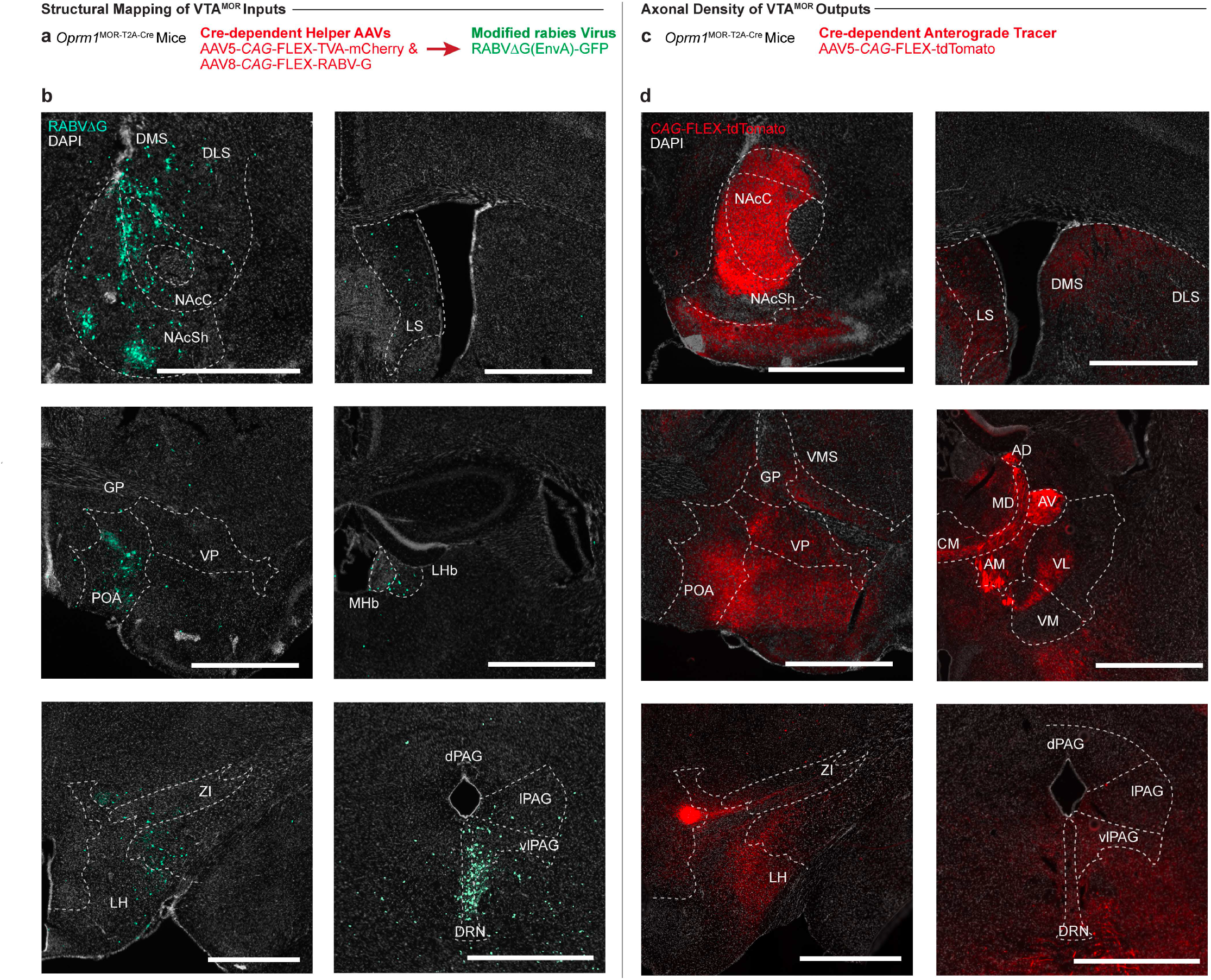
Rabies-mediated monosynaptic inputs to VTA^MOR^ neurons and anterograde tracing of VTA^MOR^ axonal outputs. **(a)** Schematic of helper AAVs and rabies virus unilaterally injected in the VTA of *Oprm1*^MOR-T2A-Cre^ mice to map **(b)** monosynaptic inputs of VTA^MOR^ neurons. **(c)** Anterograde virus unilaterally injected into the VTA of *Oprm1*^MOR-T2A-Cre^ mice to label **(d)** axonal outputs of VTA^MOR^ neurons. Scale, 1000 μm.

**Extended Data Table 4-1:**
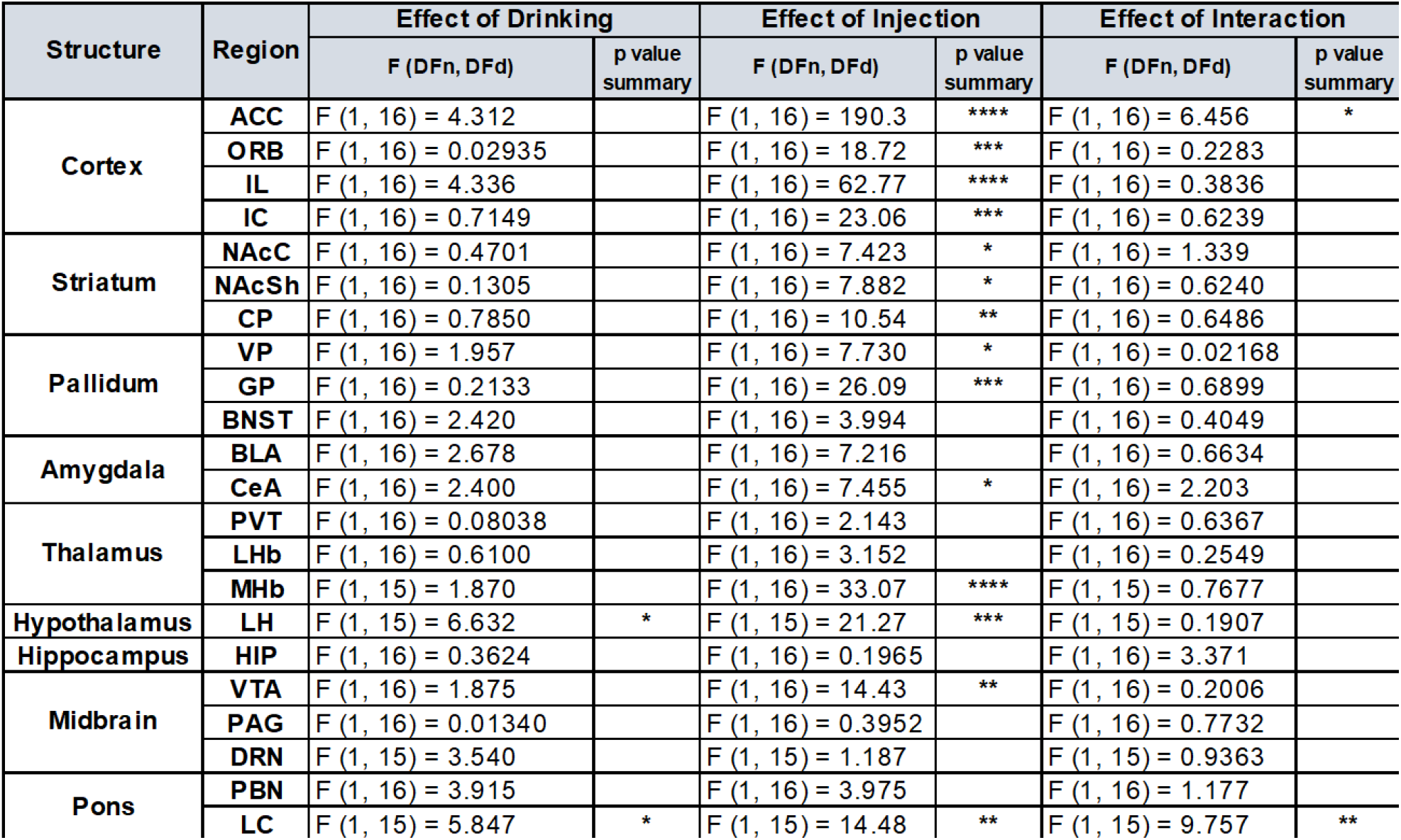
Two-Way ANOVA results for neuronal activation in response to drinking and injection in Figure 4.

**Extended Data Figure 5-1:**
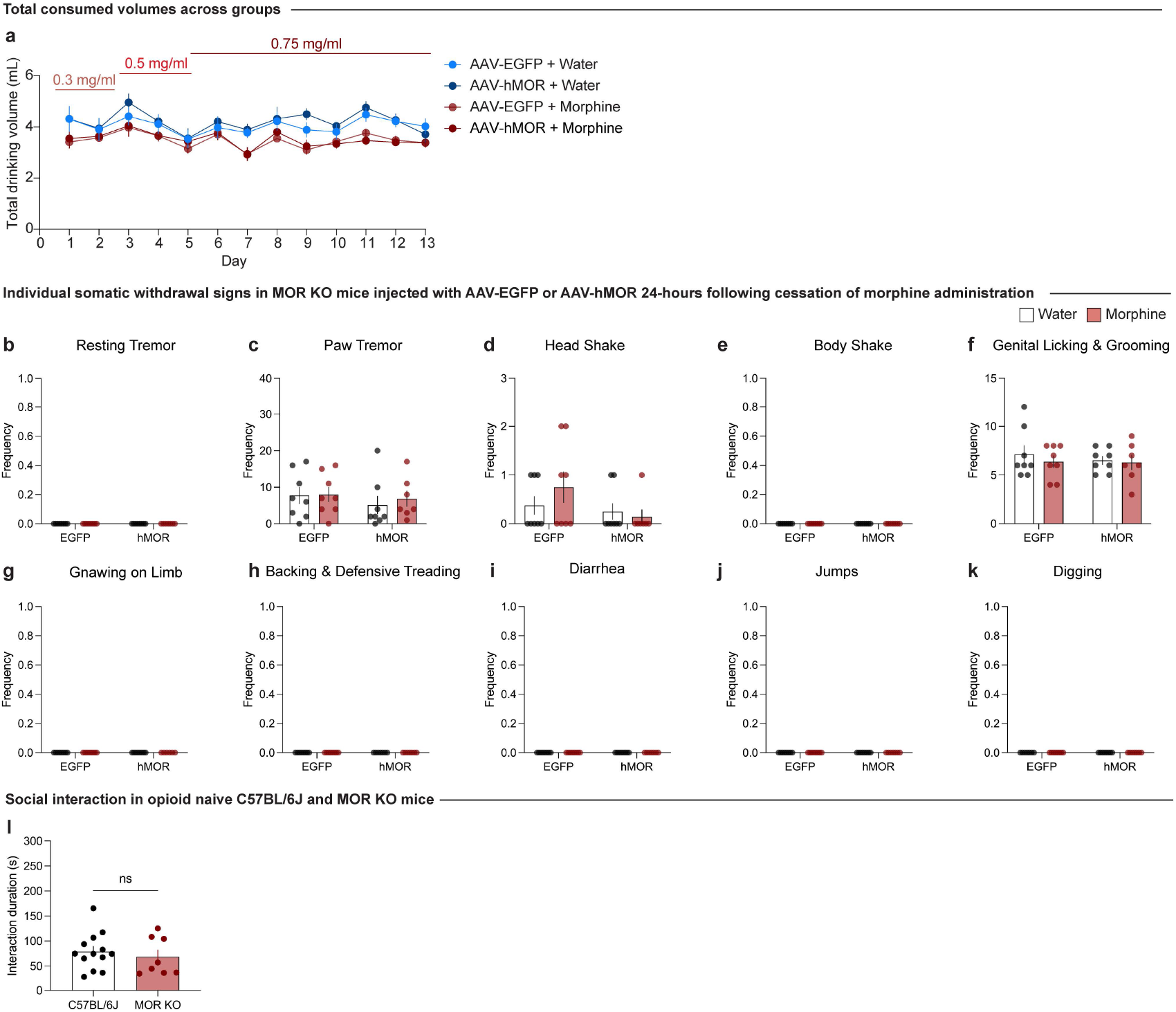
Individual somatic withdrawal signs in morphine-dependent mice undergoing 24-hour spontaneous withdrawal. (a) Total consumed drinking volume, individual somatic withdrawal signs 24-hour following cessation of morphine exposure, and social interaction in opioid naive C57BL/6J and MOR KO mice. **(a)** Total consumed volume in AAV-EGFP and AAV-hMOR injected groups exposed to water or chronic morphine show no differences in volume consumed each day compared to day 1 across all groups (two-way ANOVA + Bonferroni’s multiple comparison, day F_5.014,154.2_=7.215, p<0.0001; group F_3,31_=3.776, p=0.0203; interaction (day x group) F_36,369_=0.9681, p=0.5249). Frequencies of **(b)** resting tremors, **(c)** paw tremors, **(d)** head shakes, **(e)** body shakes, (**f)** genital licking and grooming, **(g)** gnawing on limb, **(h)** backing and defensive treading bouts, **(i)** jumps, **(j)** jumps, and **(k)** digging bouts across viral groups remain unchanged across treatment conditions. **(l)** Social interaction duration is not significantly different between opioid naïve C57BL/6J mice and MOR KO mice (injected with AAV-EGFP control virus 8 weeks prior to the social interaction test). Although these two cohorts were not conducted concurrently, all experiments were conducted under consistent conditions, including the same experimenter, facility, equipment, and handling protocols. This comparison aims to demonstrate the lack of inherent social interaction deficits in the MOR KO line used in our study (unpaired t test, t=0.6239, df=19, p=0.5401). Data are represented as mean ± SEM.

## Notes

### Competing Interest Statement

The authors have declared no competing interest.

